# Reaction-diffusion patterning of DNA-based artificial cells

**DOI:** 10.1101/2022.03.24.485404

**Authors:** Adrian Leathers, Michal Walczak, Ryan A. Brady, Assala Al Samad, Jurij Kotar, Michael J. Booth, Pietro Cicuta, Lorenzo Di Michele

**Affiliations:** Biological and Soft Systems, Cavendish Laboratory, University of Cambridge, Cambridge CB3 0HE, UK; Department of Chemistry, Faculty of Natural and Mathematical Sciences, King’s College London, London SE1 1DB, UK; Chemistry Research Laboratory, University of Oxford, Oxford OX1 3TA, UK; Department of Chemistry, University College London, London WC1H 0AJ, UK; Department of Chemistry, Imperial College London, Molecular Sciences Research Hub, London W12 0BZ, UK; fabriCELL, Imperial College London, Molecular Sciences Research Hub, London W12 0BZ, UK

## Abstract

Biological cells display complex internal architectures, with distinct micro environments that establish the chemical heterogeneity needed to sustain cellular functions. The continued efforts to create advanced cell mimics – *artificial cells* – demands strategies to construct similarly heterogeneous structures with localized functionalities. Here, we introduce a platform for constructing membrane-less artificial cells from the self-assembly of synthetic DNA nanostructures, in which internal domains can be established thanks to prescribed reaction-diffusion waves. The method, rationalized through numerical modeling, enables the formation of up to five distinct, concentric environments, in which functional moieties can be localized. As a proof-of-concept, we apply this platform to build DNA-based artificial cells in which a prototypical nucleus synthesizes fluorescent RNA aptamers, which then accumulate in a surrounding storage shell, thus demonstrating spatial segregation of functionalities reminiscent of that observed in biological cells.

## Introduction

Bottom-up synthetic biology aims to engineer artificial systems that exhibit biomimetic structure and functionality from the rational combination of molecular and nanoscale elements. These systems often take the form of *artificial cells* (ACs), micro-robots constructed *de novo* to replicate a subset of the behaviors typically associated with biological cellular life, including communication, adaptation, energy conversion, and motility.^1–3^ Despite being still far from the complexity of live cells, ACs are regarded as promising technological platforms for personalized healthcare, where cell-like microdevices could operate *in vivo,* detect disease-related biomarkers and respond by synthesizing and releasing therapeutic agents, potentially resulting in minimally toxic and efficient treatments.^1,4–7^ Similarly, ACs could underpin innovations in synthesis, through the optimized production of materials and pharmaceuticals, and in environmental remediation by selectively capturing and storing pollutants^1,4,5,8,9^.

ACs often rely on micro-compartments constructed from lipid,^10,11^ polymer^11,12^ or protein membranes,^13^ but membrane-less implementations based on coacervates^14–19^ or hydrogels^18–21^ are gaining traction, driven by enhanced robustness, easy manufacturing, and the renewed interest for biomolecular condensates and membrane-less compartments in cell biology.^22,23^ Similar to the case of biological condensates, accumulation of target molecules within membrane-less ACs can be induced through selective affinity for the scaffold phase, without relying on a membrane.

Like (eukaryotic) cells, AC implementations can benefit from an heterogeneous internal architecture to regulate transport and spatial distribution of (bio)molecules, which can facilitate the design of biomimetic pathways requiring the co-localization or separation of specific compounds.^24^ However, while with membrane-based platforms it is relatively easy to establish internal heterogeneity, for instance through nesting or sequential assembly,^24^ no general platform has been proposed to program local composition in membrane-less scaffolds. Here, we leverage the structural and dynamic programmability afforded by DNA nanotechnology^25,26^ to construct membrane-less condensates of DNA nanostructures, which can be “patterned” thanks to a reaction-diffusion scheme^27–30^ (Fig. 1a). This strategy can generate up to five chemically addressable, distinct micro-environments in a concentric geometry, whose features can be rationalized through numerical modeling. As a proof-of-concept, we use the platform to create model ACs with spatially resolved functionality, namely where a fluorescent RNA aptamer is synthesized in a prototypical “nucleus” and accumulates in an outer shell (Fig. 1a, right).

**Figure 1:**
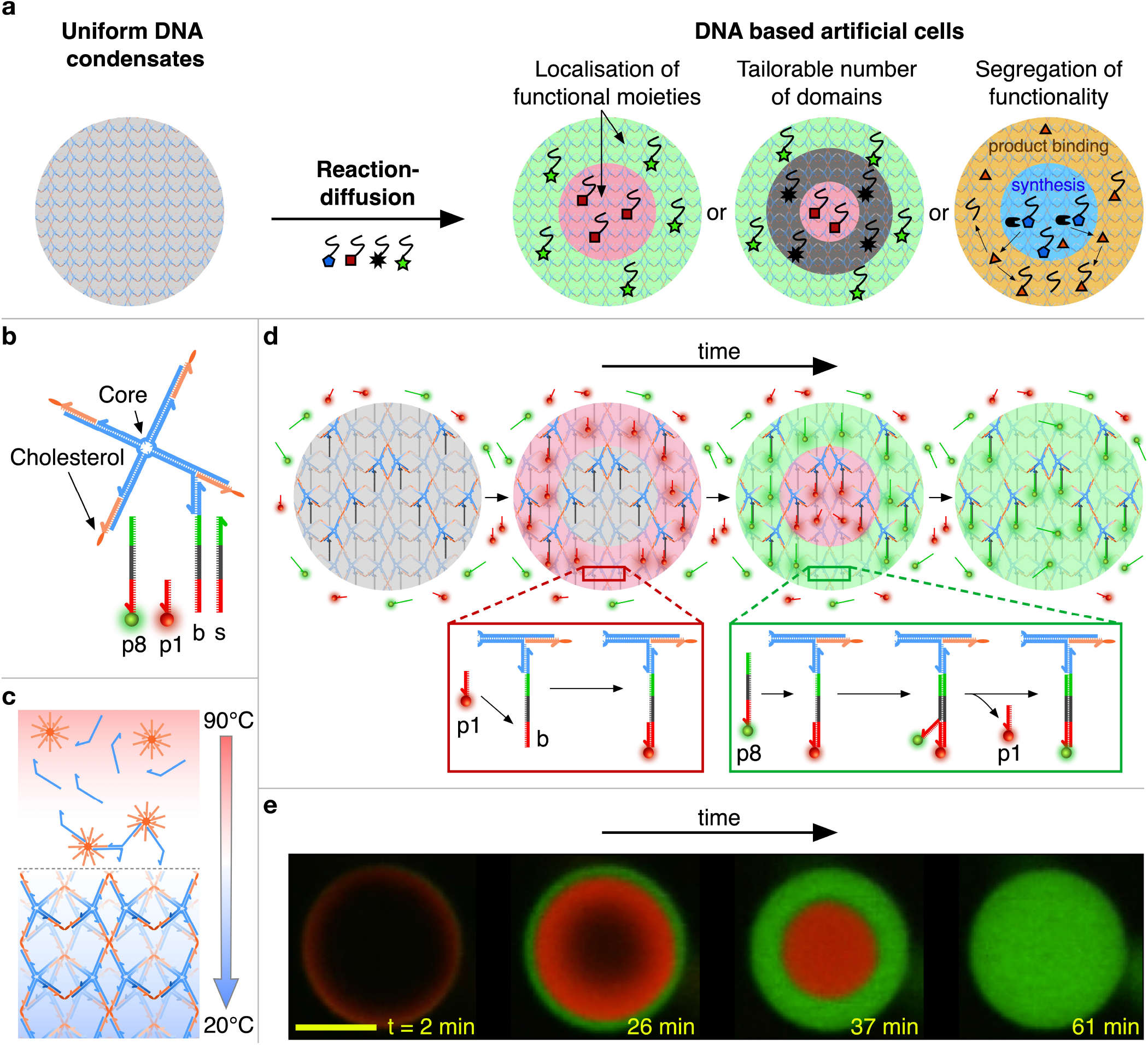
Reaction-diffusion patterning of amphiphilic DNA condensates. **a:** Reaction-diffusion processes are used to pattern initially uniform DNA condensates and construct DNA-based artificial cells featuring distinct internal environments of controllable number and molecular makeup, unlocking spatial engineering of functionality. **b:** Foundational building block of the condensates consisting of a locked four-way DNA junction with cholesterol moieties at the end of each arm. The constructs are composed of four distinct strands forming the junction (blue) and four identical cholesterolized oligonucleotides (orange).^31–33^ One arm features an additional overhang to which a *base* strand (b) is connected, which serves as a binding site for complementary, freely diffusing *patterning* strands. The latter range between 16 nt (p1) to 40 nt (p8) in length, and compete over (color-coded) overlapping binding domains on the base strand. p1 and p8 shown here are functionalized with Alexa 594 (red) and Alexa 488 (green) fluorophores, respectively. The *stop* strand (s) has the same sequence as the base and can be added in solution to sequester the excess patterning strands. Sequences of all DNA oligonucleotides are provided in Table S1, SI. **c:** Assembly process for amphiphilic DNA condensates. Samples containing all single-stranded DNA components are slowly annealed from 90°C to 20°C leading to the formation of a nano-porous framework with the branched DNA motifs connecting micelle-like hydrophobic cores where the cholesterol modifications localize, as previously reported.^31–33^ Details on sample preparation are provided in the Methods (SI). **d:** Schematic depiction of the designed reaction-diffusion pathway. At time *t* = 0, condensates are exposed to a solution of p1 (short, red) and p8 (long, green) patterning strands in excess concentration compared to base strands. Short p1 DNA strands are able to diffuse inside the condensates faster than long p8 strands, allowing for prior binding to the base strand (red box). At later times, p8 strands then diffuse within the condensates and, due to the sequence design, are able to displace p1 strands *via* toeholding^36,37^ (green box). The result is a sequence of two fronts propagating inward through the condensate. **e:** Series of confocal micrographs of the process discussed in **c**, where propagating fronts are visualized thanks to fluorescent modifications of p1 and p8. Scale bar: 15 μm.

## Results and Discussion

Our condensates self-assemble from branched, amphiphilic DNA nanostructures, shown in Fig. 1b. Similar constructs were previously demonstrated to form nanoporus phases with programmable structure, molecular-sieving properties, stimuli responsiveness, and the ability to host dynamic DNA circuitry.^31–35^ As depicted in Fig. 1c, Figure S1 (SI), and detailed in the Experimental Methods (SI), spherical aggregates with cell-like dimensions (10 - 40 μm in diameter) readily emerge from a one-pot annealing reaction. Small Angle X-Ray Scattering demonstrates that the condensates display internal crystalline order with a BCC unit cell and lattice parameter of 26.8 nm, consistently with previous observations on similar systems (Figure S2).^32^

The foundational building block used here consists of a DNA four-way junction, with cholesterol moieties labelling the end of each 35-base-pair (bp) duplex arm (Fig. 1b). One of the arms is modified with an additional single-stranded overhang to which a *base* (b) strand is connected. The base strand serves as a competitive binding site for freely diffusing *patterning strands* (p) of different lengths as depicted in Fig. 1b, where complementarity of domains is shown by same coloration and opposite directionality. All the patterning strands feature an identical (red) domain complementary to the base, but for longer strands complementarity is extended to more adjacent domains. This feature allows any longer patterning strand to displace a shorter strand from the base *via toehold mediated strand displacement* (toeholding),^36,37^ but not *vice versa,* establishing a length-dependent binding hierarchy. The length of even the shortest binding domain (14 nucleotides - nt) is such that thermal detachment of the patterning strands does not occur within experimentally-relevant timescales. Sequences of all oligonucleotides are reported in Table S1, SI.

The principle for AC patterning is schematically depicted in Fig. 1d. Condensates are prepared hosting uniformly distributed base strands but without any patterning strands initially connected. Multiple types of patterning strands are then introduced in solution, with excess concentration compared to the number of available binding sites in the condensates (see Experimental Methods, SI). In this example, we introduce patterning strands p1 (16 nt) and p8 (40 nt) labelled with an Alexa 594 (red) and Alexa 488 (green), respectively. Note that we label patterning strands with numbers increasing with strand length, such that p(n + 1) is longer than pn. The diffusion coefficient of DNA decreases with contour length,^38^ and this dependency is enhanced in porous environments like our condensates,^39^ allowing shorter DNA strands to diffuse significantly faster than longer ones. The shorter patterning strands, p1 in the example, will thus rapidly access the condensate, occupying base strands progressively from the outside of the condensate inwards. At later times, the longer p8 strand also diffuse through the condensate and, as they do so, displace p1 strands from the base strands *via* toeholding, releasing them back into solution. The result, experimentally demonstrated through confocal micrographs in Fig. 1e, is a sequence of inward-propagating fluorescent fronts: a red wave corresponding to the rapidly diffusing p1 appears first, which is then replaced by a green front produced by the slowly diffusing but strongly binding p8 strands. Note that, in confocal measurements, signal from excess patterning strands in solution is not visible due to the comparatively much higher concentration of binding sites within the condensates.

This scheme thus allows us to localize different oligonucleotides in distinct and individually addressable concentric shells within the condensates. While in this example the patterning strands bear a simple fluorescent modification, one could easily envisage including functional elements, thus paving the way for the establishment spatially resolved functionality in membrane-less ACs.

Domain structure and evolution can be programmed *via* the number and length of patterning strands, as summarized in Fig. 2. Confocal micrographs for representative condensates exposed to one, two, three, and five patterning strands are shown in Fig. 2a-d-ii while SI Videos S1-S12 exemplify pattern evolution in individual condensates and larger sample areas (see supplementary videos key, SI). In Fig. 2a-d-iii, the spatiotemporal evolution of the patterns is captured by 2D color maps showing the (azimuthally averaged and normalized) radial profile of the fluorescence intensity, *I(r,t),* where the r is the distance from the centroid of the condensate and *t* is the time elapsed from exposure to the patterning strands (see Experimental Methods). Examples of analogous color maps for multiple condensates and different numbers of patterning strands are shown in Figs S3-6 (SI). For tests with more than two domains, non-fluorescent (dark) patterning strands of lengths intermediate between two fluorescent ones were used. For instance, in Fig. 2c, a dark 30 nt strand (p6) is used in combination with 16 nt (red) p1 and 40 nt (green) p8, generating a dark shell that separates the fast-propagating red wave and the slowest-propagating green wave. Two dark (p3, p7) and three fluorescent strands (p1, p5, p8) are used in Fig. 2d, generating five distinct microenvironments at ~7 minutes from the exposure to the patterning strands, as highlighted by the azimuthally averaged fluorescent-intensity profiles (Fig. 2d-iv). Note that in this case the difference in length between adjacent species varies between 5 and 7 nt, demonstrating a separation ability comparable with electrophoretic techniques and hinting at possible applications of the ACs for the detection and separation of nucleic acids. For a fixed number of patterning strands, the relative width of the domains can be controlled by design, as shown in Fig. S7 (SI) for three-domain experiments where dark strands of different lengths are used.

**Figure 2:**
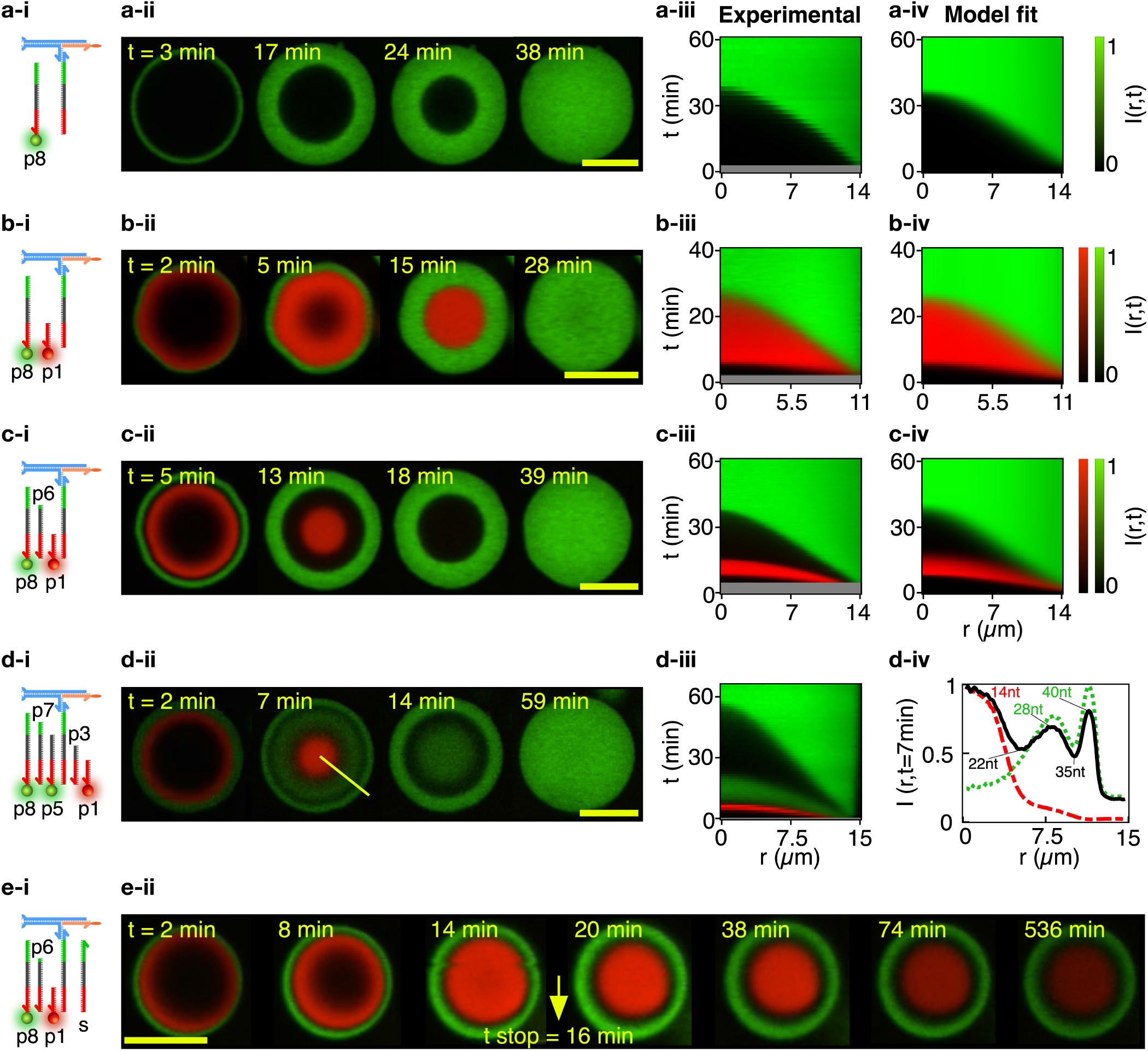
Condensate patterning is predictable and customizable. **a-d**: Patterning-strand scheme (**-i**) and equatorial confocal microscopy sections (**-ii**) for condensates patterned to form an increasing number of concentric domains, from one in **a** to five in **d**. Some patterning strands are fluorescently labeled with Alexa 594 (p1) and Alexa 488 (p5, p8), while others do not bear modifications, hence resulting in dark regions intermitting fluorescent shells in confocal data. See Table S1 (SI) for DNA sequences. The spatiotemporal evolution of the domain structure is visualized as the azimuthally-averaged, normalized radial intensity profile, *I(r,t*), where, *r* is the radial coordinate defined from the centroid of the condensate and t is the time elapsed from exposure of the condensates to the patterning strands (**-iii**). For systems **a**-**c**, I(r, t) is compared with the fitted outcome of a reaction-diffusion numerical model (**-iv**). Note that early times are not shown in experimental color maps (gray bands) due to a delay between the time at which condensates are exposed to patterning strand (t = 0) and the start of confocal recording. See Methods (SI) for information on image analysis and numerical modeling. For system **d**, sub-panel **-iv** shows radial intensity profiles extracted from confocal images at t = 7 min, highlighting the presence of five distinct domains. The green dotted and red dashed lines mark the signal from the Alexa 488 (p5, p8) and Alexa 594 (p1) channels, respectively, while the black solid line represents the overall intensity. All profiles are normalized by their highest value. **e:** Domain propagation can be arrested by adding an excess of the stop strand (s) in solution (**-i**, see also Fig. 1a), demonstrated in **-ii** with confocal data for a system with three patterning strands (p1, p6, p8). The stop strand is added at t = 16min, after which no further pattern evolution is observed (besides photobleaching). Videos S1-S8 show pattern evolution in individual condensates (even numbered) and larger fields of view (odd numbered). See supplementary videos key in SI. Scale bars: 15 μm.

In all examples discussed, the reactions are designed to progress towards the equilibrium configuration in which the longest, most stable construct occupies all binding sites, defying the purpose of our strategy as a means of engineering internal AC architecture. Pattern evolution can however be readily arrested using a *stop* strand (s, Fig. 1a), with sequence identical to that of the base. The stop strand is added in solution with excess concentration compared to that of all patterning strands combined (see Experimental Methods), sequestering them and interrupting wavefront propagation. Figure 2e-ii shows that patterns arrested with this protocol remain stable for several hours, as required for the purpose of spatial engineering in ACs. The lack of any visible blurring of the patters confirms the absence of internal diffusion of the amphiphilic DNA building blocks that make up the structure of the condensates.

The system’s evolution can be modeled through a set of coupled reaction-diffusion equations under the assumption of a spherical condensate geometry and excess patterning strands in solution, as fully detailed in the Modeling Methods (SI). Alongside known or easily determined system parameters such as condensate size, the model requires as input the diffusion constants *(D)* of the patterning strands, the second-order rate constants through which the patterning strands bind the base and/or displace previously bound strands (*k*_on_), and the exchange rate of patterning strands between the bulk and the condensate (*k*_in_).^40,41^ The latter three quantities (*D, k*_on_ and *k*_in_), graphically depicted in Fig. 3a, are used as fitting parameters. The model also features a partition coefficient of the patterning strands within the condensates^40^ that, for realistic values, is found to have no significant effect on the fitting outcomes, and was thus set to 1 (Fig. S8).

**Figure 3:**
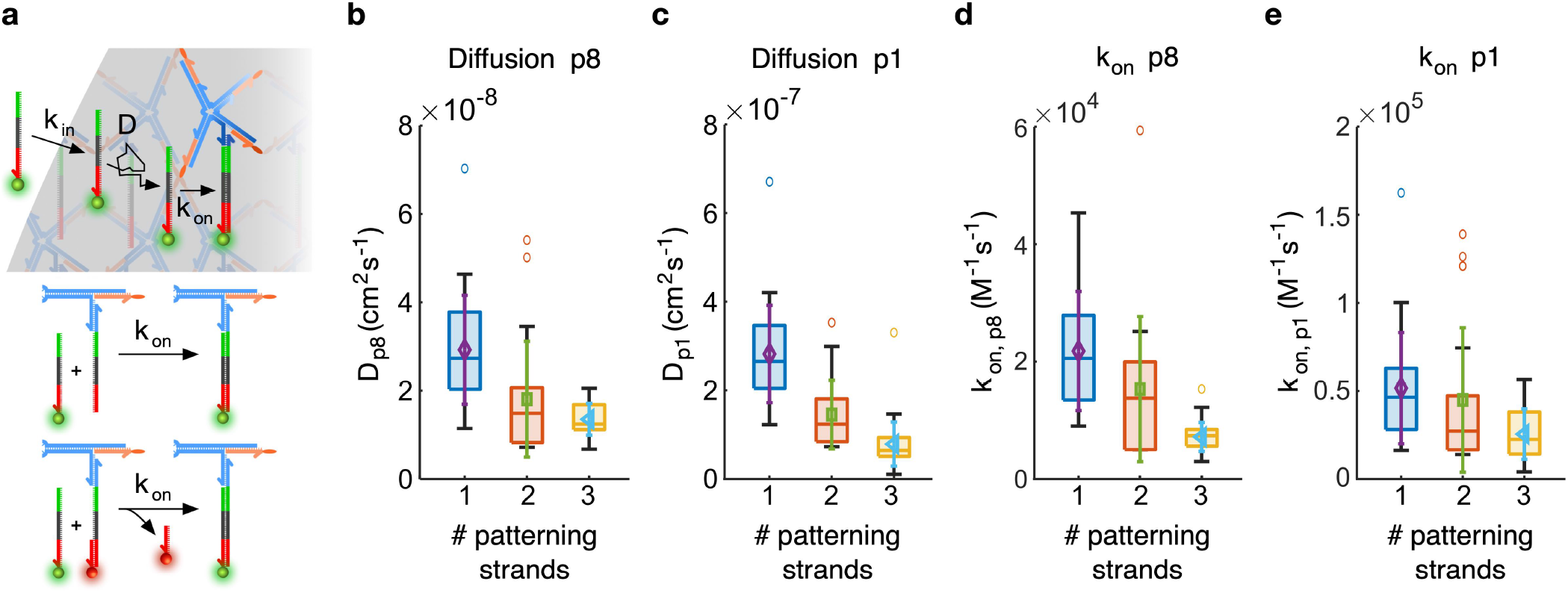
Model fitting enables extraction of reaction-diffusion parameters. **a:** Schematic representation of the color parameters consisting of an entry rate *k*_in_, a diffusion constant *D* and a binding/displacement rate *k*_on_ (top). With *k*_on_ we indicate both the second order binding rate of a patterning strand to a free binding site and that of the toehold-mediated strand displacement process through which a longer patterning strand replaces a shorter one previously occupying a binding site (bottom, see Modeling Methods). **b-c:** Diffusion coefficients for 40 nt patterning strand p8 (**b**) and 16 nt patterning strand p1 (**c**). **d-e:** Binding rates for 40 nt patterning strand p8 (**d**) and 16 nt patterning strand p1 (**e**). Data are shown for samples with one patterning strand (p1 or p8; Fig. 2a and Figs S3-4; *N* = 33 condensates for p1 and *N* = 29 condensates for p8), two patterning strands (p1 and p8; Fig. 2b and Fig. S5; N = 23 condensates) and three patterning strands (p1, p6, and p8; Fig. 2c and Fig. S6; N = 43 condensates). The results are displayed as box plots with highlighted median, upper and lower quartiles (box), and 50th centiles (whiskers) outliers excluded. Overlaid to the box plots are the mean (symbol) and standard deviation (same-color errorbar) of the distributions.

The model outputs the spatiotemporal evolution of the concentration of bound patterning strands within the condensates, which, after accounting for diffraction-induced blurring and normalizing, renders an estimate of the experimental fluorescent intensities. This model can thus be used to fit the experimental I(r, t) data for up to three competing patterning strands (see Modeling Methods). Qualitative comparison between experimental and fitted I(r, t) maps demonstrates good agreement, as visible in Fig. 2a-c-iv and Figs S3-6. A quantitative assessment of the fit residuals, shown in Fig. S9, confirms a good match, with deviations typically within ±10 — 15% for data associated to long (p8) patterning strands and within ±20 — 25% for short (p1) strands. The larger deviation observed for shorter patterning strands is ascribable to the smaller datasets, given that short strands experience a briefer reaction-diffusion transient. In Fig. S10 we further show histograms of the residuals combining values from all sampled condensates. The distributions appear quite symmetrical but deviate from a normal profile. A non-normal distribution may result from the the small shape differences between modeled and experimental reaction-diffusion profiles, also highlighted in the residual maps (Fig. S9). These differences may emerge due to early-time effects related to sample mixing, optical artifacts due to refractive-index mismatches and absorption, non-spherical condensate geometry, or nonisotropic diffusion caused by the contact with the bottom of the experimental chamber.

Figure 3b-c summarizes the distributions of fitted diffusion constants and binding coefficients for our shortest (p1, 16 nt) and longest (p8, 40 nt) patterning strands, as determined for experiments with a single patterning strand (p1 or p8; Fig. 2a and Figs S3-4), two patterning strands (p1 and p8; Fig. 2b and Fig. S5), and three patterning strands (p1, p6, and p8; Fig. 2c and Fig. S6).

We observe an order-of-magnitude difference in diffusion constant between p1 and p8, with *D*_p1_ = 1–4× 10^-7^ cm^2^ s^-1^ and *D*_p8_ = 1–4× 10^-8^ cm^2^ s^-1^. *D* is the primary parameter that determines the propagation speed of the reactiondiffusion fronts through the condensates, therefore, the difference found between *D*_p1_ and *D*_p8_ is consistent with expectations. We further note that both diffusion constants decrease in the presence of additional patterning strands, hinting at crowding effects.

Figure 3d-e shows the fitting outcomes for rate constant *k*_on_. In all cases, *k*_on, p1_ describes binding of p1 to a free base strand, given that p1 is unable to displace other incumbent strands. Similarly, *k*_on, p8_ describes hybridization to free base strands in experiments only featuring p8. When shorter patterning strands are present alongside p8, *k*_on, p8_ can be interpreted as the effective second-order rate constant of the toehold-mediated strand displacement reaction through which p8 displaces incumbent p1 (2-strand case) and p6 (3-strand case).^36^ See Fig. 3a (bottom), and Modeling Methods for a quantitative justification of this interpretation. Fits produce values of *k*_on, p1_ and *k*_on, p8_ within the same order of magnitude (10^4^ – 10^5^ M^-1^ s^-1^), with the former slightly larger than the latter. Values are consistent with previous observations for hybridization in hydrogels,^28^ and smaller than rates typically measured in freely diffusing constructs (*r*~ 10^6^ M^-1^ s^-1^).^36^ The similarity between rates of hybridization (*k*_on, p1_) and toeholding (*k*_on, p8_) is expected, given that the two are known to converge for toehold lengths in excess of 6 nt - a condition that is verified in our reactions.^36^ A decreasing trend is found with increasing number of patterning strands, which is more pronounced for *k*_on, p8_ compared to *k*_on, p1_ and may thus be due to crowding.

The material exchange rate *k*_in_, as summarized in Fig. S11, is slightly larger for p1, compared with p8, consistent with the higher diffusion rate of the shorter patterning strand, while no significant trend is observed as a function of the number of patterning strands.

To explore possible correlations between the fitting parameters and gauge the robustness of our fits we computed maps of χ^2^ as a function of each pair of parameters (Fig. S12).^42–44^ The maps reveal a degree of correlation between D and *k*_in_, while *k*_on_ is uncorrelated to both other variables. All maps, however, show a clear minimum hinting at identifiability of all parameters, which is confirmed by the individual likelihood curves and associated identifiability analysis (Fig. S13). We note that, while *k*_on_ is identifiable, the model displays relatively weak sensitivity to this parameter for values in excess of ~ 10^5^ M^-1^ s^-1^ (Figs S12a and S13a). This weakened sensitivity can be rationalized as the result of the diffraction blurring applied in the model, which for sufficiently large *k*_on_ dominates over the blurring of the propagating front induced by finite reaction rates (see Modeling Methods for details).

Having identified a route for establishing addressable domains in condensates, we can proceed with localizing functional elements within different regions to create a model AC hosting a spatially organized pathway. As summarized in Fig 4a, the AC has been patterned with a “nucleus” region hosting a *template* construct (t), containing a T7 promoter sequence and the DNA complementary sequence to an RNA aptamer, and the shell region containing binding sites for the RNA product. The template is anchored to the base strand through a *bridging* (r) strand, which ensures the formation of the double-stranded T7 promoter required for the T7 polymerase to begin transcription.^45^ The RNA produced is a modified version of the DFHBI-binding fluorescent Broccoli aptamer.^46^ Addition of an extra 8 bps to the stem region produced a significantly brighter aptamer as discussed in the Experimental Methods and Fig. S14. The aptamer also features a binding site complementary to the single-stranded overhang present in the *capture* (c) strands located in the shell region. Note that because the b-t complex located in the nucleus is longer than the capture strand present in the shell, patterning of these devices needed to be conducted following multi-step protocol, detailed in the Experimental Methods and Fig. S15 (SI). The protocol also involves washing steps to remove any unbound b-t complexes that would result in RNA synthesis occurring in the bulk solution.

**Figure 4:**
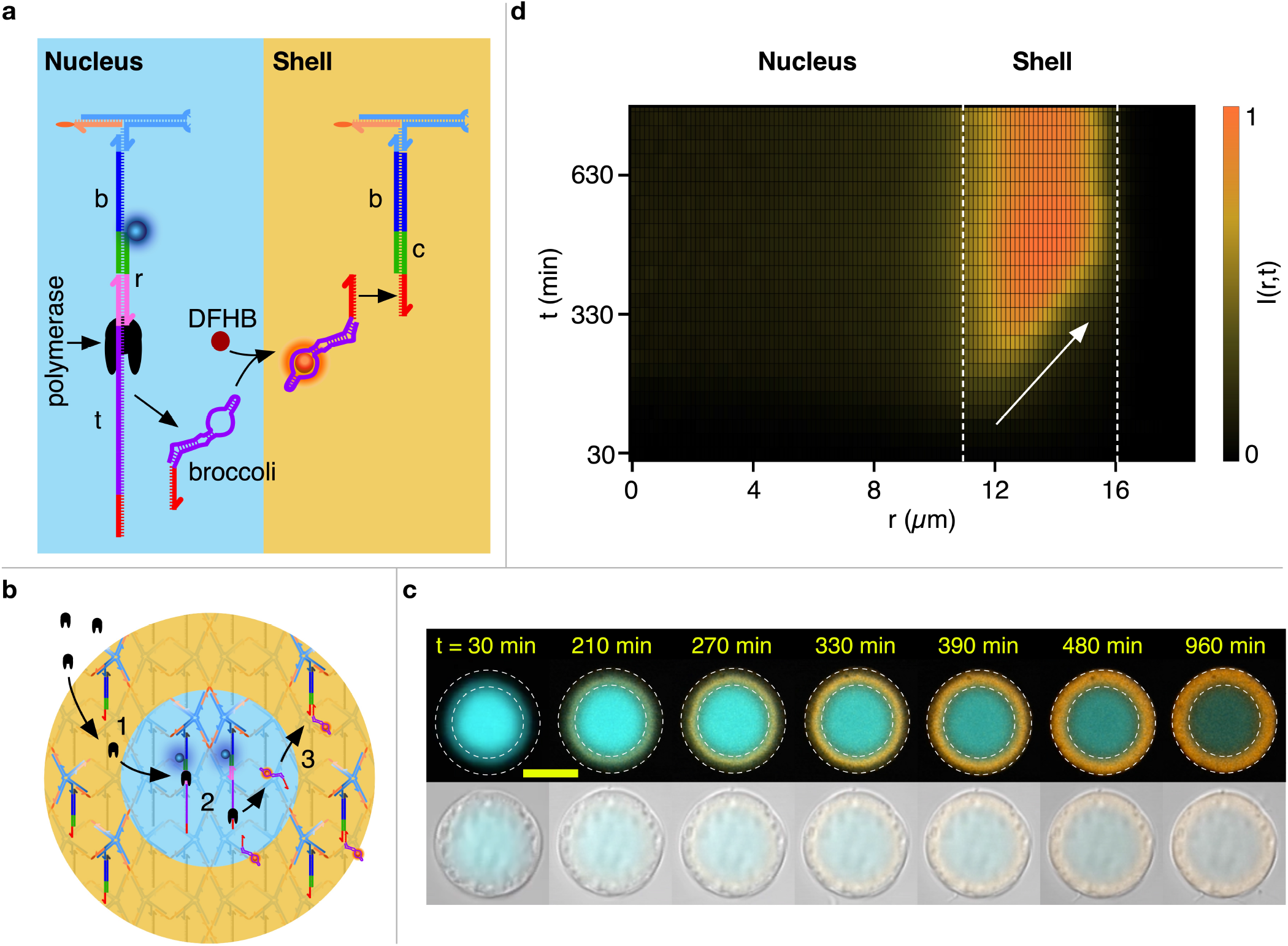
Spatially-distributed functionality in a model artificial cell. **a:** Schematics of the functional nucleic acid machinery in *nucleus* (cyan) and *shell* regions (orange). In the nucleus, connected to the base strand are a *bridge* (r) strand and the *template* (t) strand. Together, these form a double stranded T7 promoter (pink) and a single stranded polymerase template (purple, red) from which a polymerase (black) is able to synthesize Broccoli RNA aptamers (folded purple and red) that form a complex with DFHBI molecules to become fluorescent (orange). The base strands in the shell region are connected to *capture* strands (c) with single stranded overhangs (red) complementary to a free domain on the broccoli aptamer. Complementary DNA/RNA domains are shown in the same color. Protocols for patterning the ACs are detailed in the Experimental Methods (SI). **b:** Mode of operation of the AC. The polymerase is added in solution alongside NTPs, DFHBI, and other components required for Broccoli synthesis, which diffuse through the shell (1) reaching the nucleus, where the aptamers are produced (2). The aptamers then diffuse outwards and bind to the dedicated sites in the shell (3). **c:** A series of confocal images (top) of an AC progressively building up Broccoli aptamer in the shell (orange). Note how the signal accumulates from the nucleus-shell interface and propagates outwards. The nucleus is shown in cyan, and progressively photobleaches. The dashed lines mark the physical boundary of the AC and that between the nucleus and the shell. Bottom: bright field images of the same AC overlaid to (faint) confocal data, demonstrating that no physical change to the AC occurs during Broccoli synthesis. The reaction is initiated at time t = 0, as discussed in the Experimental Methods (SI). **d:** color map showing the evolution of the radial fluorescent intensity of the aptamer. The slope of the fluorescent front signals accumulation from the inside-out, as highlighted by the white arrow. Dashed lines mark the nucleus/shell and shell/solution boundaries. Videos S13-S16 show the response of multiple ACs with different shell/nucleus size ratios. Scale bar: 15 μm.

The sought response at the AC level is sketched in Fig. 4b. Patterned ACs are exposed to polymerase that, similar to other proteins of comparable size,^32^ can diffuse through the condensates, reaching the template in the nucleus. Here, the Broccoli aptamer is produced, which readily binds to DFHBI, diffuses out towards the shell and binds the capture motifs. These constructs would thus display the envisaged separation of functionality, as well as a basic form of communication between two domains, one producing a signal in the form of RNA constructs and the other receiving it.

Figure 4c (top) shows a time-resolved sequence of confocal micrographs from an AC producing the designed response. The nucleus (cyan), fluorescently stained thanks to a fluorophore on the bridging strand, retains the same size through the experiment and only undergoes progressive bleaching. In turn, fluorescence from the Broccoli aptamer (orange) builds up in the shell region from the inside-out, consistent with the RNA product being produced in the nucleus. Combined bright field / confocal micrographs of the same objects confirm that the overall size and appearance of the condensate does not change during Broccoli build up (Fig. 4c, bottom). In Fig. 4d we show a color map of the radial intensity of the Broccoli emission versus time, where the outwards-propagating front is clearly observable, marked by an arrow. Data from more ACs with different shell thicknesses relative to nucleus size are summarized in Fig. S16, while time lapses of the process can be inspected in SI Videos S13-S16. As a control, in Fig. S17 we show the results of experiments with template/promoter constructs free in solution, which expectedly show outside-in accumulation. Finally, in Figs S18 and S19 we report the time-dependent Broccoli emission in bulk fluorimetry experiments, including samples with aptamer-expressing ACs and samples that only contain the supernatant solution but no ACs. The lack of signal from the latter further confirms that aptamer production occurs exclusively within the AC nucleus.

## Conclusion

In summary, we have introduced a general process for the creation of stable and individually addressable domains in condensates selfassembled from DNA nanostructures. The process relies on a reaction-diffusion scheme, where patterning constructs with different diffusivities and binding affinities compete for binding sites within the condensates. The number and size of the domains can be tuned by design and predicted by numerical modeling. As a proof-of-concept, we adapt the patterning scheme to construct a model DNA-based artificial cell with an active nucleus that produces a fluorescent RNA aptamer, and a storage shell where the product progressively accumulates. With this basic implementation as a starting point, one could envision future developments towards artificial cells capable of producing, storing and later releasing therapeutic RNA elements, such as small-interfering RNAs.^47^ These implementations, however, would need to encapsulate also the RNA polymerase and NTPs rather than relying on freely-diffusing enzymes and building blocks, which could be achieved by surrounding the artificial cell with a less permeable barrier.

More generally, we argue that reaction-diffusion patterned DNA condensates could constitute a versatile platform for engineering cell-like agents with spatially- and temporally-resolved functionality. Potential responses are not limited to expression or capture of RNA products, as the domains can be enriched with virtually any functional molecule or nanoscale agent that can be linked to the patterning strands, including enzymes, nanoparticles, aptamers and photo-responsive elements, thus unlocking the opportunity to engineer ever more complex reaction pathways within the ACs. Finally, while here DNA condensates are used as passive scaffolds, one can envisage design modifications where patterning alters the local physical structure. For instance, one could ensure that localization of responsive moieties makes specific regions in the condensates sensitive to targeted degradation (photo-induced or enzymatic), enabling the creation of voids within the artificial cells that can be used to store bulky cargoes or simply regulate the connectivity of the remaining domains. In turn, dynamic changes in the local mesh size could enable the design of pathways that couple changes in local transport properties to biochemical activity, giving rise to more complex time-dependent responses. For example, one could envisage a negative-feedback loop where local accumulation of an RNA product represses transcription by (sterically) hindering polymerase diffusion and configurational freedom. If coupled with enzymatic RNA degradation, similar systems could sustain fluctuations in RNA expression periodic in time and space, reminiscent of oscillating gene expression in biological cells.^48^ Similarly, RNA expression could be coupled to changes in artificial cell size, leading to mechanical actuation, useful to engineer propulsion or shape changes in artificial tissues.

Acknowledgement LDM acknowledges support from a Royal Society University Research Fellowship (UF160152) and from the European Research Council (ERC) under the Horizon 2020 Research and Innovation Programme (ERC-STG No 851667 – NANOCELL). AL and LDM acknowledge support from a Royal Society Research Grant for Research Fellows (RGF/R1/180043). MJB is supported by a Royal Society University Research Fellowship (URF/R1/180172). MJB and AAS acknowledge funding from a Royal Society Enhancement Award (RGF/EA/181009) and an EPSRC New Investigator Award (EP/V030434/1). MW acknowledges support from the Engineering and Physical Sciences Research Council (EPSRC), and the Department of Physics at the University of Cambridge (the McLatchie Trust fund). The authors acknowledge Diamond Light Source for provision of synchrotron beamtime (SM28071) and thank A. Smith for assistance in operating beamline I22.

A data set associated to this publication is available free of charge at https://doi.org/10.17863/CAM.87545.

## Supporting Information Available

The Supporting Information is available free of charge at [LINK].

All experimental, image analysis, numerical modeling and fitting methods. Supplementary Figures S1-S19 including bright field microscopy images of the condensates (Fig. S1), SAXS characterization of the condensates (Fig. S2), additional experimental and fitted *I(r,t)* patterning maps (Figs S3-S6), data demonstrating control over domain thickness (Fig. S7), analysis of the fitting model behavior, correlation between fitting parameters and their idenfitiability (Figs S8-S13), performance of the extended Broccoli aptamer (Fig. S14), patterning protocol for RNA-synthesizing ACs (Fig. S15), additional data on RNA synthesis in ACs (Fig. S16), control experiments on RNA-synthesizing ACs (Figs S17-S19). Supplementary Table S1 showing nucleotide sequences. Description of Supplementary Videos S1-S16 (PDF).

Supplementary Videos S1-S16 showing condensate patterning experiments (Video S1-S12) and RNA-synthesis by ACs (Videos S13-S16) (avi).

