## Supporting Information for "Reaction-diffusion patterning of DNA-based artificial cells"

### Experimental methods

#### Strand design

Strand sequences were designed with NUPACK[1] not allowing more than three consecutive A,C,T or G nucleotides, other than where was necessary due to sequence specificity of the Broccoli aptamer. Strands were bought from Integrated DNA Technologies (IDT) including chemical modifications with Cholesterol, Alexa 488, Alexa 594 and Alexa 647. Non modified strands were purified by the supplier using standard desalting, except for the Broccoli template strand, which was purified with PAGE purification. Chemically modified strands were purified by the supplier with HPLC purification. All strands were reconstituted in 1× Tris-EDTA (TE) buffer (10mM Tris, 1 mM EDTA, pH 8.0, Sigma Aldrich) upon receipt. The concentration of the reconstituted DNA strands was determined by absorbance at 260 nm with a Nanodrop 2000 spectrophotometer (Thermo Scientific). Extinction coefficients for each strand were provided by the supplier. All sequences are listed in Table S1.

#### Saline buffers

All the following components used for saline buffers such as NaCl, $MgCl_2$, KCl, TE and Tris-HCl were purchased from Sigma Aldrich and dissolved or diluted to desired concentrations in milliQ water. Prior to use, all saline buffers were filtered through 0.22 µm pore size Millipore syringe filters (Millex). For condensate assembly, reaction diffusion experiments, washing steps and patterning processes, a saline buffer containing 1× TE and 300 mM NaCl was used. For RNA transcription experiments, whether in the bulk or within the artificial cells, a saline buffer containing 50 mM $MgCl_2$, 50 mM KCl, 40 mM Tris-HCl (pH 8.0), 2 mM Spermidine (New England Biolabs), 10 mM DTT (New England Biolabs) was used. This saline buffer is based on the RNAPol buffer by New England biolabs at 1× concentration, but with added salt.

#### Condensate assembly

All condensates were fabricated in rectangular Glass capillaries of dimension 60 mm × 4 mm × 0.4 mm (CM Scientific) as a one pot self-assembly process, adapted from Brady *et al.*[2,3] All the DNA components that make up the amphiphilic DNA nanostars (core1-4 and cholesterol in Table S1), and the base strand were mixed together in stoichiometric ratio for a target concentration of 7 µM of the nanostars (7 µM of each core strand, 7 µM of base strand and

28 µM of cholesterol strand), for a total volume of 60 µl in an Eppendorf tube. The mixture was then transferred into a capillary with a pipette. Finally, both ends of the capillary were capped with equal amounts of mineral oil and glued shut with epoxy glue (Araldite Rapid). Capillaries were then subjected to a thermal annealing process in a tailor-made, Peltier-coltrolled water bath. The samples were heated up to 90°C, incubated for 30 min, then cooled from 90°C to 63°C at a rate of -0.01°C, and finally from 63°C to 20°C at -0.1°C. The thermal annealing protocol has been modified with respect to Brady *et al.*,[2,3] to favor the emergence of spherical condensates rather than faceted crystallites that form at slower cooling rates. Note however that the condensates retain crystalline microstructure (Fig. S2). To extract the formed condensates, both ends of the capillary were cut open with a diamond tip pen. The capillary was then placed vertically in a 1.5 ml Eppendorf tube with additional 60 µl of saline buffer and spun at low rotation on a tabletop centrifuge for about 30 s. The final concentration of nanostars that make up the extracted condensates was thus diluted to 3.5 µM.

#### Condensate washing

Samples were washed by adding 150 µl of saline buffer to the Eppendorf tube containing the condensates. The sample was spun down on a table top centrifuge until the condensates reach the bottom of the tube, then 150 µl are removed from the supernatant. This process was repeated five times every time a sample was being washed. Washing was performed to remove excess components that may be present due to uncertainties in concentration measurements and pipetting or to remove components when these are used in excess during patterning steps. This is particularly critical for RNA expression experiments in which excess bridge/template can lead to Broccoli synthesis in solution, rather than only within the artificial cell nucleus.

#### Patterning of ACs for RNA expression experiments

DNA condensates for reaction-diffusion experiments required no additional preparation steps other than washing, prior to experiments. For RNA expression experiments, the condensates needed to be patterned with the bridge (r), template (t) and capture (c) strands to generate the sought core-shell architecture prior to RNA expression experiments (see Fig. 4). This was done in a multi-step process, as shown in Fig. S15:

1. Freshly extracted condensates were washed, as described above (see Fig. S15a).

2. Pre annealed bridge-template (b-t) constructs (see Table S1 and Fig. 4) were added to the condensates at a target concentration of 10 µM ($\sim 3\times$ excess), and left to diffuse inside the condensates and bind the base strands over 36 hours, ensuring full saturation of the available binding sites. At this stage, the condensates were uniformly functionalised with bridge-template constructs able to express Broccoli in the presence of T7 polymerase – the ACs were all-nucleus (see Fig. S15b).

3. The sample was washed repeating the steps outlined above, and may have been separated into aliquots to be "patterned" for different times and create shells of different thicknesses (see Fig. S15c).

4. Capture strands (see Table S1 and Fig. 4) were added at a target concentration of 15 µM and left to displace the bridge-template constructs for a desired amount of time. With reference to Fig. 4 and Fig. S15d, note that the capture (c) strand displaces the bridge (r) with a strongly thermodynamically driven toehold-mediated strand displacement reaction.

5. To arrest the patterning process, the stop2 strand (see Table S1) was added at a target concentration of 30 µM to the sample. The stop2 strand sequesters all the freely diffusing capture strands, but does not displace those already bound to the base strand as it lacks a toehold for this displacement reaction. Note that the stop2

strand, different from the stop (s) strand used in patterning experiment outlined in Fig. 2, also has partial complementarity with the bridge strand, and thus was able to separate freely diffusing bridge-template constructs (released in step 4) without forming a full length double stranded promoter region for the T7 RNA polymerase. This precautionary measure was designed to deactivate displaced promoter-template constructs whilst keeping the ones bound to the core of the condensate intact, and was aimed at further ensuring that the only viable templates for the polymerase were within the AC nucleus (see Fig. S15e).

6. Finally, the condensates were washed again (see Fig. S15f).

While we opted for the multi step strand displacement process outlined above, one could achieve the same patterning for RNA-expressing ACs also by simultaneously introducing b (not previously hybridized to t) and c strands, as these two follow the established diffusion/binding hierarchy rules explored in Fig. 2 (*i.e.* b diffuses faster than c, but c displaces b). After the addition of the stop strand, this process would produce condensates with a b-rich core and a c-rich shell. To reach the final product, however, an additional step would still be needed where the long t strand is introduced and left to hybridize with the b strands in the core. Given the similar experimental complexity of the two approaches, we ultimately opted for the method described in Fig. S15 as it allowed us to pre-anneal the b-t complex, hence ensuring defect-free hybridization. The absence of defects in the b-t constructs is indeed critical as the b-t duplex region functions as promoter region for the T7 RNA polymerase, whose recognition ability can be compromised by the presence of defects.

#### Confocal microscopy

Confocal microscopy images and videos were obtained using a Leica TCS SP5 scanning laser confocal microscope with a HC PL APO CORR CS 40×/0.85 dry objective (Leica). Samples were illuminated by either Ar-ion laser lines (454 nm, 488 nm) or He-Ne laser lines (594 nm,

633 nm) depending on the fluorophore of interest. Alexa 488 was excited at 488 nm, Alexa 594 at 594 nm, Alexa 647 at 633 nm and the Broccoli-DFHBI complex at 454 nm. The fluorescent signal was recorded using the Leica Photomultiplier (PMT) detectors. Acquisition windows were set at $500 - 580$ nm for Alexa 488, $610 - 750$ nm for Alexa 594, $645 - 750$ nm for Alexa 633 and $480 - 580$ nm for for the Broccoli-DFHBI complex. Brightfield images were recorded at the same time using the transmission photomultiplier detector (Fig. S1). All condensates were imaged as timelapse sequences of z-stacks at hand picked positions within the chambers. The acquisition process was automated using the default Leica software. RNA expression experiments in artificial cells were conducted at 37°C thanks to a an environmental microscopy chamber.

#### Reaction-diffusion patterning experiments

Reaction-diffusion experiment samples were prepared by pre-mixing the desired patterning strand species at a target concentration of 4 µM with saline buffer (1×TE, 300 mM NaCl) in a solid quartz microscopy multi-wells slide (Hamamatsu). The sample was then mounted on the microscopy stage and secured with Blu Tack. Condensates were then added to the mixture at a target concentration of base strands (or nanostars) equal to 0.4 µM. The dense condensates rapidly started sinking towards the bottom of the cell, while the diffusion-reaction process was initiated. The chamber was quickly sealed with FlexWell sealing strips (Sigma Aldrich) to prevent evaporation. Imaging positions were then quickly selected and the experiment launched, trying to minimize the time elapsed from insertion of the condensates to the start of recording. This time was recorded with a stopwatch and taken into consideration during analysis (note gray bands in the experimental color-maps, Fig. 2). Note that flow experienced by the condensates as they sink may affect the reaction-diffusion process at the earliest stage.

Experiments in which the stop strand was added were conducted in the same way but with a reduce patterning-strand concentration of 2 µM to slightly slow down the reaction-diffusion process and allow for a longer delay between the start of the experiment and the addition of the stop strand. After the desired wait, the stop strand was added at a final concentration of two times the concentration of patterning strand and species (4 µM for 1 patterning strand, 8 µM for 2 strands, etc.). For these experiments, the top of the well was only sealed after the addition of stop strand.

#### RNA expression in artificial cells - microscopy experiments

RNA expression experiments in the artificial cells were undertaken in a similar fashion as the patterning experiments. Here, the polymerase saline buffer was loaded in a well with 40 mM of each NTP (ATP, CTP, GTP, UTP) (Thermo Fisher Scientific), 60 µM DFHBI, 0.042 µM of T7 RNA Polymerase (New England Biolabs). The artificial cells (condensates pre-patterned as discussed above) were included at a target nanostar (and thus template) concentration of 0.35 µM . The well plate was sealed with FlexWell sealing strips and imaged at 37°C using the (pre-heated) environmental chamber.

#### Image analysis

The confocal microscopy time series of z-stacks were analyzed using custom software written in Matlab to obtain the intensity maps $I(r,t)$, shown in the main text (Figs. 2, 4) and SI (Figs S3-S7). Leica files (.lif) were imported using the Bio-Formats toolkit and then sorted according to imaging position, z stack plane and acquisition time. Condensates were selected for analysis to be as spherical as possible (ensuring accurate comparison with the model) and not in contact with other condensates, which could render patterning strand diffusion anisotropic. For each condensate, we first identified the confocal slice closest to the equatorial plane, which was done manually, and then subtracted the background signal measured by pixel average over manually selected rectangular regions that did not contain any condensates. Let us refer to time series of background-subtracted equatorial images as $J(x, y; t)$, where $x$ and $y$ are the Cartesian (discrete) pixel coordinates and $t$ is the time elapsed from exposure of the condensates to the patterning strands. We then proceeded to localizing the centroid of the condensate, which was done following a hybrid manual/automated approach, where the approximate centroid position was at first selected by hand *via* an interactive user interface, and then refined through an image segmentation pipeline comprising of the following steps: contrast enhancement, binarization, dilation, erosion and finally *imfindcircle* – a circle finding function. The latter step produced an accurate location of the centroid, which was cross-referenced with the manually selected centroid positions. Condensates showing a poor match between manually selected and refined centroid positions were discarded. Images were processed in reverse chronological order because the condensates were fully fluorescent and easily recognizable at the end of the experiment but dark at the start. Pixels in $J(x, y; t)$ were then binned based on the distance $r$ from the centroid. The average pixel intensity for each radial-distance bin was then extracted, which is analogous to carrying out an azimuthal average of the intensity in a polar coordinate system. This angular average was calculated for each time-point to obtain a 2D intensity map, $J(r, t)$. The final $I(r, t)$ maps were calculated by normalizing – $I(r, t) = J(r, t)/\max(J(r, t))$ – so that $0 \leq I(r, t) \leq 1$. For experiments where data were recorded in both green and red channels, $I(r, t)$ was computed for both channels. In these experiments, centroid localization was only conducted for the channel corresponding to the longest patterning strand, and then used to compute $I(r, t)$ in all channels.

#### RNA expression – bulk fluorimetry experiments

Bulk fluorimetry experiments were conducted to probe Broccoli synthesis in control experiments, both from free templates and artificial cells (Figs S18-S19), using a FluoStar Omega plate reader with a 450 nm excitation filter and a 520 nm emission filter. Samples with the polymerase saline buffer as described above were loaded in 384-well plates (Greiner 384, flat bottom). Strand concentrations used are the same as for the confocal microscopy experiments.

#### Characterisation of extended vs original (short) Broccoli aptamer

Data for the fluorescence of the short (original) and extended (used in this work) Broccoli aptamer, shown in Fig. S14, were obtained as follows. Short or Extended Broccoli template strands and a T7 promoter strand were mixed in an equimolar ratio in milliQ water. The strands were heated to 95°C for 1 min and then slowly cooled to 25°C at a rate of -0.1°C $s^{-1}$. The annealing reactions were held for 10 s at 25°C before cooling down to 8°C. In vitro transcription reactions were prepared on ice with a total reaction volume of 10 µl. The final concentrations of the reaction components was 100 ng template per 10 µl reaction, 1 mM Tris pH 7.4, 2 mM spermidine, 45 mM $MgCl_2$, 10 mM NTPs each, 1 mM DTT, 50 mM KCl, 1 µl of T7 RNA polymerase (NEB), and 60 µM DFHBI. The mixtures were incubated in the PCR machine for 4 h at 37°C, 5 µl of the solutions were diluted by adding 36 µl of 10 mM tris pH 8 on ice. The fluorescence measurement of 40 µl of each mixture was quantified on a Tecan infinite M1000PRO fluorimeter (455 nm excitation, 506 nm emission, and optimal gain).

#### Small Angle X-Ray Scattering

Small Angle X-Ray Scattering measurements and data evaluation were undertaken according to the methodology of Brady *et al*.[2,4] DNA condensates produced as described above were transferred into X-ray capillaries and left to sink until forming a pellet at the bottom. Powder diffraction patterns were acquired from this pellet at the I22 beamline of Diamond Light Source and the resulting diffraction patterns were processed and fitted with custom Matlab software, as described in Brady *et al*.[2,4] Data are shown in Fig. S2 and discussed in the caption.

### Modeling methods

#### Reaction-diffusion model

The reaction-diffusion processes inside condensates were modeled with the equations of Fickian diffusion with additional terms describing binding of the patterning strands to the base strands. The condensates were assumed to be porous, perfect spheres with immobile and evenly distributed binding sites, allowing for patterning strands to diffuse throughout the condensates and interact with the binding sites. For the diffusion aspect, the spherical assumption allows for reduction of the three dimensional diffusion equation to a one dimensional equation in radial coordinates.

We modeled hybridization of the patterning strands to free binding sites as an irreversible second order process, neglecting the possible presence of short-lived partial binding events, which would not be detectable in confocal microscopy experiments. Binding irreversibility over the experimental timescales is guaranteed by a complementarity of at least 14 bp between patterning and base strands (see Table S1). The following chemical equation was thus used to describe patterning-strand hybridization in our model

$$b + p_i \xrightarrow{k_{\mathrm{on},i}} b\,p_i, \qquad \text{(S1)}$$

where $b$ indicates a *free* binding site (base strand), $p_i$ is the patterning strand (p1-p8 in the notation of the main text), $k_{\mathrm{on},i}$ is the binding rate, and $b\,p_i$ is the bound state of the patterning strand to the base strand.

Combining the above assumptions for diffusion and binding we obtain the following system of partial differential equations (PDE) for a single patterning strand species

$$\begin{cases} \frac{\partial [p_i]}{\partial t} = D_i \frac{1}{r^2}\frac{\partial}{\partial r}(r^2 \frac{\partial}{\partial r}[p_i]) - k_{\mathrm{on},i}\,[b][p_i] \\ \frac{\partial [b]}{\partial t} = -k_{\mathrm{on},i}\,[b][p_i] \\ \frac{\partial [b\,p_i]}{\partial t} = k_{\mathrm{on},i}[b][p_i], \end{cases} \tag{S2}$$

where $D_i$ is the diffusion constant of the patterning strand, $r$ is the radial coordinate, $t$ is time, and $[\dots]$ indicates concentrations. Due to the assumption of immovable binding sites, $[b\,p_i] = [b_0] - [b]$ can be substituted in the third line of Eq S2, where $[b_0]$ is the initial concentration of free binding sites in the condensates.

Toehold-mediated strand displacement, through which a patterning strand displaces another from the base strand, was also modeled as an irreversible second-order reaction, neglecting transitional binding states. This assumption is justified because, in all cases, the length of the toehold is at least six nucleotides, leading to fast equilibration toward the product and short lived transitional binding states.[5] The resulting chemical equation is

$$b\,p_i + p_j \xrightarrow{k_{\mathrm{on},ij}} b\,p_j + p_i, \tag{S3}$$

where $p_i$ is the incumbent patterning strand, $p_j$ a longer invading patterning strand and $k_{\mathrm{on},ij}$ is an effective second-order toehold-mediated strand displacement rate.

In a system with two competing patterning strands, the differential equation system then becomes

$$\begin{cases} \frac{\partial [p_i]}{\partial t} = D_i \frac{1}{r^2}\frac{\partial}{\partial r}(r^2 \frac{\partial}{\partial r}[p_i]) - k_{\text{on},i}[p_i][b] + k_{\text{on},ij}[p_j][b\,p_i] \\ \frac{\partial [p_j]}{\partial t} = D_j \frac{1}{r^2}\frac{\partial}{\partial r}(r^2 \frac{\partial}{\partial r}[p_j]) - k_{\text{on},j}[p_j][b] - k_{\text{on},ij}[p_j][b\,p_i] \\ \frac{\partial [b]}{\partial t} = -k_{\text{on},i}[p_i][b] - k_{\text{on},j}[p_j][b] \\ \frac{\partial [b\,p_i]}{\partial t} = k_{\text{on},i}[p_i][b] - k_{\text{on},ij}[p_j][b\,p_i] \\ [b\,p_j] = [b_0] - [b] - [b\,p_i]. \end{cases} \tag{S4}$$

For simplicity, in Eq S4 we assumed that the binding rate of the longer ($j$) patterning strand, $k_{\text{on},j}$, is equal to the toeholding rate $k_{\text{on},ij}$. This assumption is justified both because the two rates are expected to be similar, and because the process of $p_j$ binding the base strand by displacing $p_i$ is expected to be strongly dominant over the process of $p_j$ binding an empty site, given that the shorter strand will have occupied nearly all binding sites by the time the longer strand reaches a given location. In other words, because $D_i > D_j$, for a given $r$, $[p_j]$ only starts increasing from 0 when $[b] \ll [b\,p_i]$, and hence the third term in the right-hand-side of the second line of Eq S4 is going to always dominate over the second term.

The model for three strands can be extended in the same way, including patterning strand $p_l$ able to displace both $p_i$ and $p_j$

$$\begin{cases} \frac{\partial [p_i]}{\partial t} = D_i \frac{1}{r^2}\frac{\partial}{\partial r}(r^2 \frac{\partial}{\partial r}[p_i]) - k_{\text{on},i}[p_i][b] + k_{\text{on},ij}[p_j][b\,p_i] + k_{\text{on},il}[p_l][b\,p_i] \\ \frac{\partial [p_j]}{\partial t} = D_j \frac{1}{r^2}\frac{\partial}{\partial r}(r^2 \frac{\partial}{\partial r}[p_j]) - k_{\text{on},j}[p_j][b] - k_{\text{on},ij}[p_j][b\,p_i] + k_{\text{on},jl}[p_l][b\,p_j] \\ \frac{\partial [p_l]}{\partial t} = D_l \frac{1}{r^2}\frac{\partial}{\partial r}(r^2 \frac{\partial}{\partial r}[p_l]) - k_{\text{on},l}[p_l][b] - k_{\text{on},il}[p_l][b\,p_i] - k_{\text{on},jl}[p_l][b\,p_j] \\ \frac{\partial [b]}{\partial t} = -k_{\text{on},i}[p_i][b] - k_{\text{on},j}[p_j][b] - k_{\text{on},l}[p_l][b] \\ \frac{\partial [b\,p_i]}{\partial t} = k_{\text{on},i}[p_i][b] - k_{\text{on},ij}[p_j][b\,p_i] - k_{\text{on},il}[p_l][b\,p_i] \\ \frac{\partial [b\,p_j]}{\partial t} = k_{\text{on},j}[p_j][b] + k_{\text{on},ij}[p_j][b\,p_i] - k_{\text{on},jl}[p_l][b\,p_j] \\ [b\,p_l] = [b_0] - [b] - [b\,p_i] - [b\,p_j], \end{cases} \tag{S5}$$

where, as above, we use $k_{\text{on},j} = k_{\text{on},ij}$ and $k_{\text{on},l} = k_{\text{on},il} = k_{\text{on},jl}$.

To describe the initial and boundary conditions of the model we use the notation $[p_i(r,t)]$, $[b(r,t)]$ and $[b\,p_i(r,t)]$ to indicate the concentration of strand $p_i$, free binding sites and binding sites occupied by $p_i$ (respectively), at the radial position $r$ at time $t$. The radial coordinate is bounded by $0 \leq r \leq R$, where R is the radius of the spherical condensate. We assumed that initially there are no patterning strands inside the condensate and no occupied binding sites, thus $[p_i(r,0)] = [b\,p_i(r,0)] = 0$ and $[b(r,0)] = [b_0]$. The total concentration of binding sites $[b_0]$ was assumed to be $r$-independent (uniform condensate composition) and estimated assuming that each nanostar within the condensate carries one binding site and occupies a volume equal to the sphere with radius given by the nanostar arm length, $n = 35\,\text{bp}$

$$[b_0] = \frac{3}{4\pi(na_0)^3 N_A} = 235.25\,\mu\text{M}, \tag{S6}$$

where $\mathrm{a_0} = 0.34$ nm is the average length of an individual nucleotide and $\mathrm{N_A}$ is Avogadro's number.

Following,[6] for the patterning strands, the boundary conditions at the outer edge ($R = r$) of the condensates are chosen as those of a permeable interface in a two-compartment system comprising of the bulk and the interior of the condensates, where binding is neglected. At the center of the condensates ($r = 0$) the boundary is considered an impermeable surface.

$$\begin{cases} \frac{\partial[p_i(r=0,t)]}{\partial r} = 0 \\ \frac{\partial[p_i(r=R,t)]}{\partial t} = k_{\text{in},i}\left([p_{i,\,bulk}] - \frac{1}{\lambda}[p_i(r=R,t)]\right) \\ [b(r=R,t)] = 0 \end{cases} \tag{S7}$$

where $[p_{i,\,bulk}] = 4\,\mu\text{M}$ is the experimentally defined bulk concentration of pattering strand $p_i$, $\lambda$ a partition coefficient for the strand $p_i$ in the condensate and $k_{\text{in},j}$ a rate describing exchange of patterning strands between the condensate and bulk. $[p_{i,\,bulk}]$ was assumed constant in the limit of excess patterning strands. Analogous initial and boundary conditions were applied to all patterning strands, if multiple ones were present. In our case, for the numerical implementation of the model the partition coefficient is taken to be $\lambda = 1$.

##### Numerical implementation

The system of PDE's describing the reaction diffusion process in the DNA condensates was solved numerically using a finite differences method described by Crank.[7] This method reduces the system of PDE's to a system of ordinary differential equations (ODE) by discretizing the radial axis into equidistant points of coordinates $r_n = n\,\delta r$ for $n = 0, 1, 2, ..., N$, where $\delta r$ is the spacing. The second order spatial derivatives can now be approximated with centered finite difference approximations in spherical coordinates, leading to a system of ODE's at every coordinate point, shown in eqs. S8-S10.

$$\frac{\partial [p_i]_n}{\partial t} = \begin{cases} k_{in}([p_{i,\,bulk}] - [p_i]_n) \quad \text{for} \quad n = N, \\ \frac{6D_i}{(\delta r)^2}([p_i]_1 - [p_i]_0) + \kappa_{i,n} \quad \text{for} \quad n = 0, \\ \frac{D_i}{n(\delta r)^2}\left\{(n+1)[p_i]_{n+1} - 2n[p_i]_n + (n-1)[p_i]_{n-1}\right\} + \kappa_{i,n} \quad \text{otherwise,} \end{cases} \tag{S8}$$

where $[p_i]_n$ is the concentration of patterning strand $p_{\text{i}}$ at position $n$, and $\kappa_{i,n}$ is the sum of binding and unbinding terms for patterning strand $p_{\text{i}}$ at position $n$. For example, in the two strand case as in eq. S4, we have $\kappa_{i,n} = -k_{\text{on},i}[p_i]_n[b]_n + k_{\text{on},ij}[p_j]_n[b\,p_i]_n$.

$$\frac{\partial [b]_n}{\partial t} = \begin{cases} \kappa_{b,n} \quad \text{for} \quad n \neq N, \\ 0 \quad \text{for} \quad n = N, \end{cases} \tag{S9}$$

where $[b]_n$ is the concentration of the binding site at position n and $\kappa_{b,n}$ is the sum of binding and unbinding terms for the binding site at position $n$.

$$\frac{\partial [b\,p_i]_n}{\partial t} = \begin{cases} \kappa_{bp_i,n} \quad \text{for} \quad n \neq N, \\ 0 \quad \text{for} \quad n = N, \end{cases} \tag{S10}$$

where $[b\,p_i]_n$ is the concentration of the patterning strand $p_i$ bound to the binding site $b$ at position $n$ and $\kappa_{\mathrm{bp_i,n}}$ is the sum of binding and unbinding terms for that strand combination at position $n$.

#### Data fitting

The numerical solution of the ODE system detailed in Eq. S8-S10, obtained using Matlab's *ode15s* differential equation solver, was used to fit experimental data. The numerical outcome consists in the (discrete) time- and radius- dependent concentrations of the bound patterning strands, $[bp_i](r,t)$. In the fitting routine, the solutions for $[bp_i](r,t)$ were additionally modified to account for finite imaging resolution and other experimental effects, as follows. First, the concentrations profiles were interpolated onto a circle of radius $R$, and convolved with a 2D Gaussian Point Spread Function (PSF) to account for experimental diffraction blurring. The width $\sigma$ for the PSF was estimated from the full width at half maximum (FWHM) of an Airy disk

$$FWHM = 0.51\frac{\lambda}{NA}, \tag{S11}$$

where $\lambda$ is the emission wavelength of the fluorophore and $NA = 0.85$ is the numerical aperture of the objective, and $\sigma$ can be calculated as

$$\sigma = \frac{FWHM}{2\sqrt{2\ln(2)}}. \tag{S12}$$

A different value of $\sigma$ was used to blur the numerical patterns depending on the fluorophore associated to the patterning strand whose concentration one wishes to fit. In the case of our fluorophores, namely Alexa 488 and Alexa 594 the wavelength at the emission peaks are 520 nm and 617 nm respectively leading to $\sigma_{520nm} = 132.5\,\mathrm{nm}$ and $\sigma_{617nm} = 157.2\,\mathrm{nm}$.

The blurred two-dimensional patterns were then once more converted to radial profiles. This profiles were additionally normalized to an experimental fluorescence-intensity radial profile acquired for the specific condensate that one intended to fit, once the condensate was fully saturated with a patterning strand. This normalization was applied to account for off-plane fluorescence projections and other possible optical effects such as those induced by refractive index mismatches and absorption.

The resulting (convolved and normalized) time-dependent radial profiles of the bound-strand concentrations are assumed to correspond directly to the normalized fluorescence intensity profiles $I(r,t)$ computed as discussed above (*e.g.* color-maps shown in Fig. 2, 4 and Figs. S3-S6), and were thus used to fit experimental data. Note that the significantly (about 2 orders of magnitude) higher concentration of binding sites in the condensates compared to the concentration of free strands in solution guarantees that the contribution from free strands is negligible in experiments, and can thus be disregarded also in the fitting model. The Matlab least squares optimization routine *lsqcurvefit* was used for the fitting.

For samples with multiple patterning strands the multi-strand numerical model was used, but rather than optimizing for the concentration profiles of all strands at once, fitting was performed sequentially for individual (fluorescent) strands, starting from the longest (slowest-diffusing) strand and then progressing to the shorter strands. When fitting the shorter strands, the parameters obtained from the previous fitting performed for the longer strands were kept constant. For instance, for a two-strand experiment with patterning strands p1 and p8, as shown in Fig. 2b, we would first optimize for concentration profile of p8, obtaining $D_{\mathrm{p8}}$, $k_{\mathrm{on,p8}}$ and $k_{\mathrm{in,p8}}$. Then we would keep these fixed and fit for the p1 profile, obtaining $D_{\mathrm{p1}}$, $k_{\mathrm{on,p1}}$ and $k_{\mathrm{in,p1}}$. This sequential fitting protocol was required because the significantly different timescales over which the multiple strands diffuse through the condensates lead to imprecise fitting for the fast-diffusing strands. Indeed, the fluorescence profiles relative to the short strands reach the center of the condensate rapidly compared to the timescales required for the long strands to complete the process, and then experience progressive photobleaching. Therefore, if one were to fit long and short strands simultaneously, for the latter the results would be heavily influenced by the long-time portion of the data which carries bleaching artifacts and little information about the reaction-diffusion kinetics, reducing the accuracy of the fitting. In turn, when fitting for one patterning strand at a time, we could simply crop the experimental data in time, focusing on the timescales relevant to each patterning strand.

For experiments featuring non-fluorescent patterning strands, fitting was first carried for all fluorescent strands sequentially, as discussed above, thus determining the corresponding rate constant and diffusion coefficients. A global fit over all fluorescent profiles was then conducted, optimizing against the parameters associated to the dark strands, while keeping the previously fitted parameters for the fluorescent strands fixed.

Initial values for the fitting parameters $D$ and $k_{\text{in}}$ were estimated by manual tweaking, bringing the numerical outcomes close to the experimental patterns. Instead, $k_{\text{on}}$ was purposefully initialized at values well below (by one order of magnitude) the values typically observed at convergence. This choice was motivated by the evidence that the numerical model becomes insensitive to the value of $k_{\text{on}}$, when this becomes too large ($\sim 10^6$), which is in turn due to the Gaussian blurring applied to account for finite imaging resolution. Indeed, the primary effect of $k_{\text{on}}$ is that of regulating the sharpness of the propagating wavefront. For large $k_{\text{on}}$, the contribution from this parameter becomes negligible compared to the applied Gaussian blurring, hence the model looses sensitivity. By initializing $k_{\text{on}}$ from low values, we ensure convergence to either a meaningful outcome or saturation to a value where the model becomes insensitive. Due to this issue, we stress that the data shown in Fig. 3d and e should be considered with caution, as they may represent lower bounds to the actual binding/toeholding rate constants.

### Supplementary figures

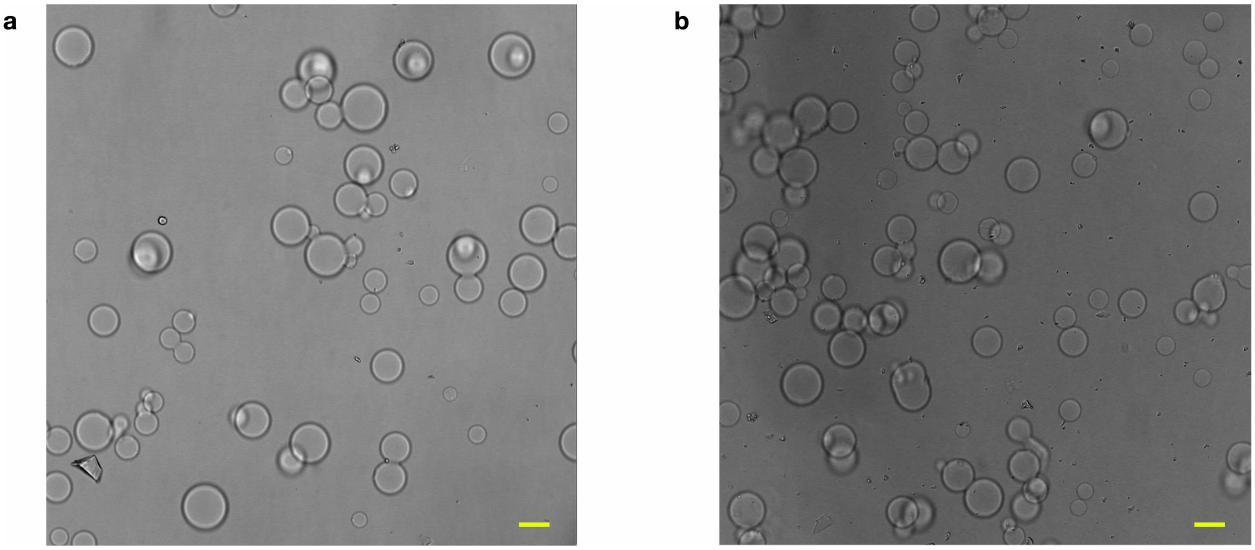

**Figure S1:** Bright field microscopy images of condensates in experimental wells. **a:** and **b:** show two fields of view. The scale bars correspond to 15 µm.

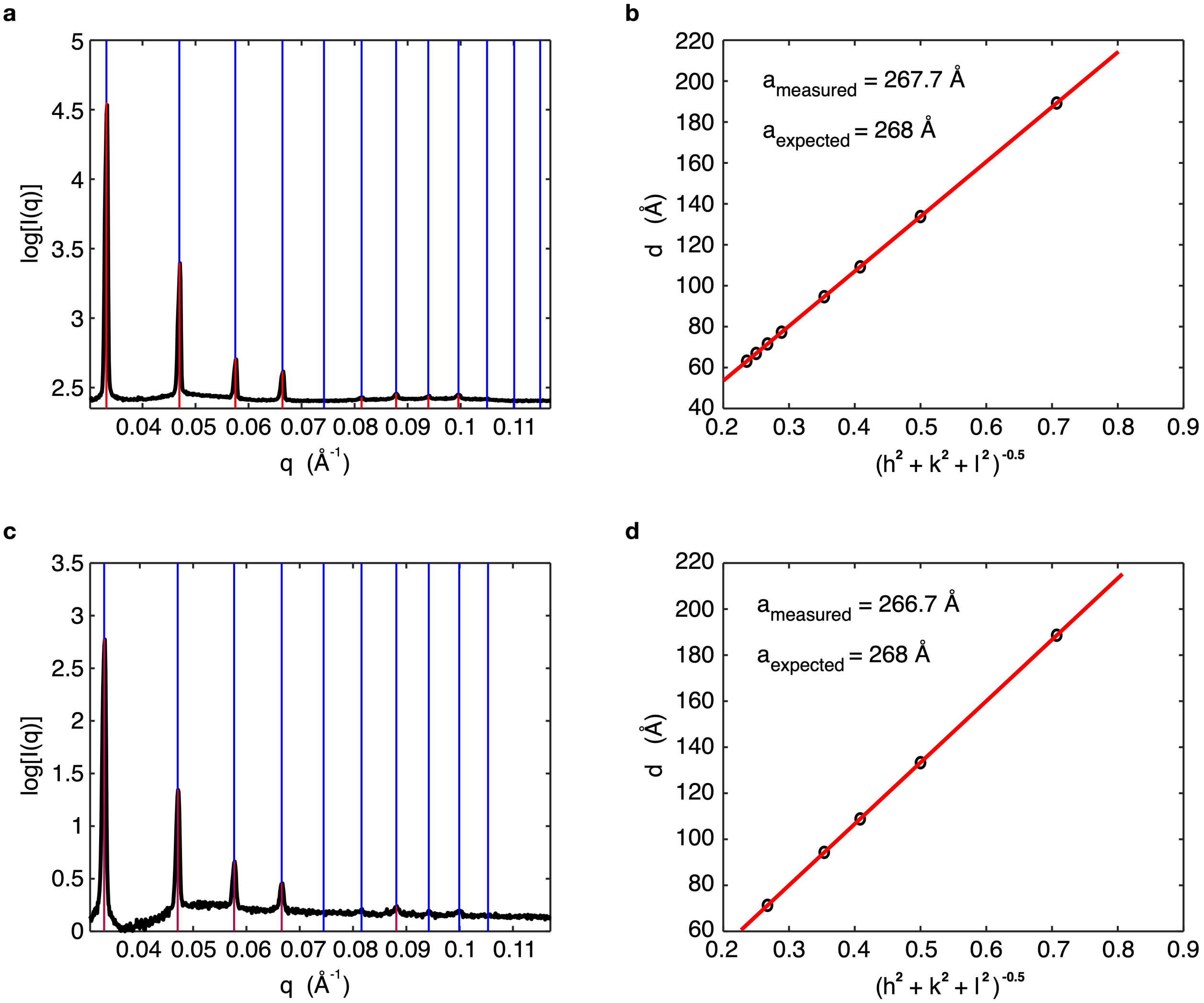

**Figure S2:** Small Angle X-ray Scattering reveals crystalline order within the condensates. **a, c**: Radial average of powder diffraction pattern acquired by exposing several condensates (pelleted in an X-ray capillary). Bragg peaks, marked by red lines, reveal crystalline order. Blue lines indicate best fits of the peaks to the expected locations for a BCC lattice, confirming the structure previously determined from assemblies of amphiphilic DNA nanostars similar to the ones used in this work.[3] **b, d**: $d$-spacing of the identified peaks as a function of $\sqrt{h^2+k^2+l^2}$, where $h$, $k$ and $l$ are Miller indices. The linear trend confirms a cubic unit cell and allows estimation of the lattice parameters. Data in **a** and **b** are relative to condensates without any strands bound to the overhang of the nanostars, while those in **c** and **d** correspond to condensates where the base strand is bound. The estimated lattice parameters $a_{\text{measured}} = 26.77\,\text{nm}$, and $26.67\,\text{nm}$ match the expected value ($a_{\text{expected}} = 26.8\,\text{nm}$) for amphiphilic DNA nanostars with arm length of 35 bp (used here), as determined by Brady *et al.* [3] The results from both data sets show no apparent structural difference in the condensates when additional strands are bound to the overhang of the nanostar.

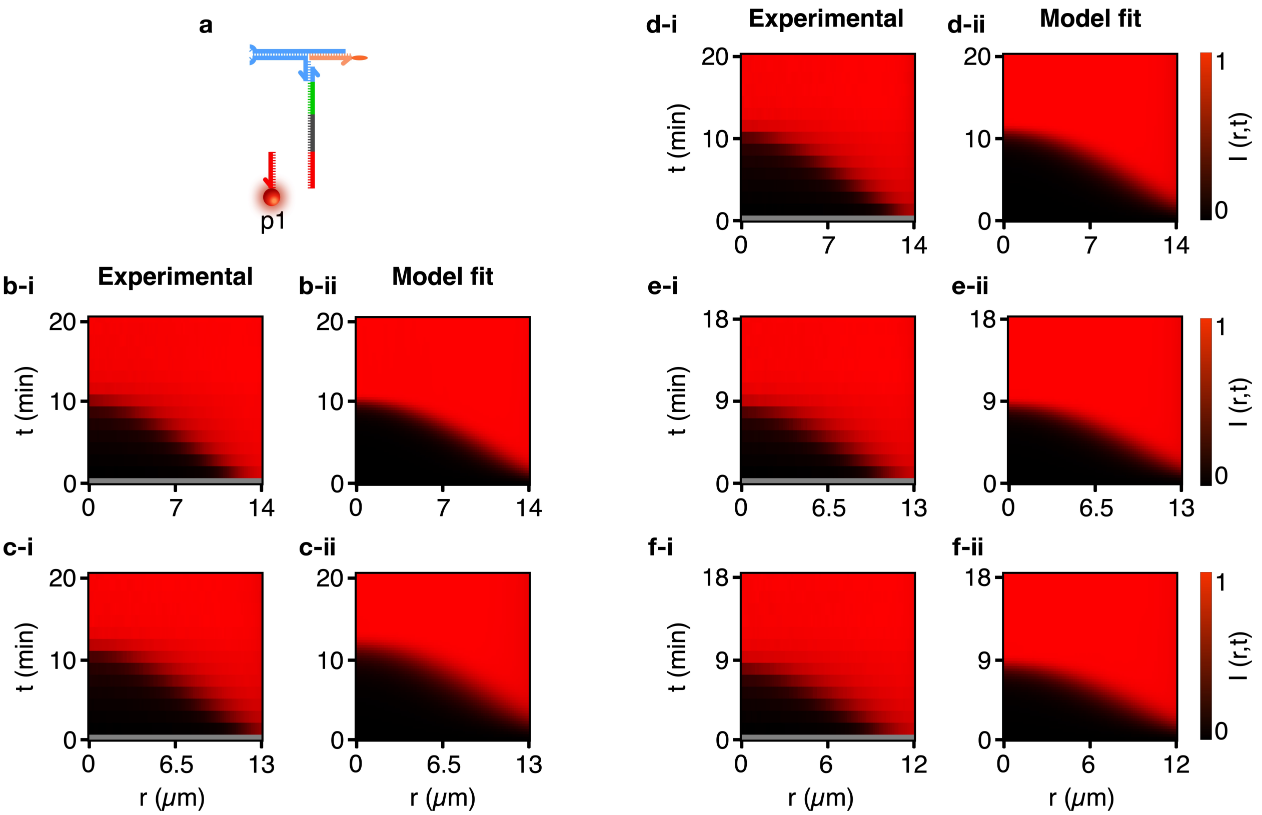

**Figure S3:** Experimental and modeled (fitted) results in 5 different DNA condensates of varying sizes showing the reaction diffusion behavior of a single patterning strand, namely p1. **a**: Schematic depiction of the patterning and base strands. **b-f**: Experimental (**-i**) and modeled (**-ii**) color maps as used and discussed in Fig. 2. Gray bands in the experimental data mark the time gap between exposure of the condensates to patterning strands and the beginning of data acquisition (see Experimental Methods).

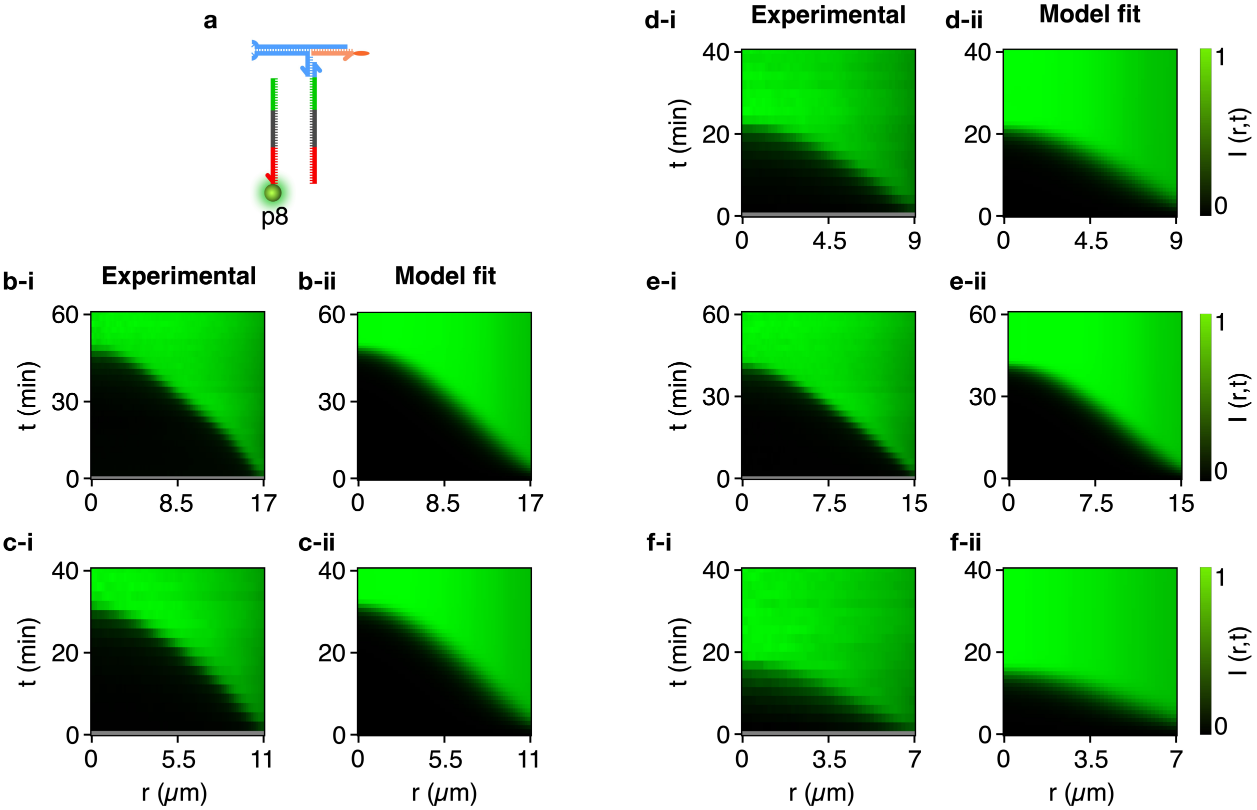

**Figure S4:** Experimental and modeled (fitted) results in 5 different DNA condensates of varying sizes showing the reaction diffusion behavior of a single patterning strand, namely p8. **a**: Schematic depiction of the patterning and base strands. **b-f**: Experimental (**-i**) and modeled (**-ii**) color maps as used and discussed in Fig. 2. Gray bands in the experimental data mark the time gap between exposure of the condensates to patterning strands and the beginning of data acquisition (see Experimental Methods).

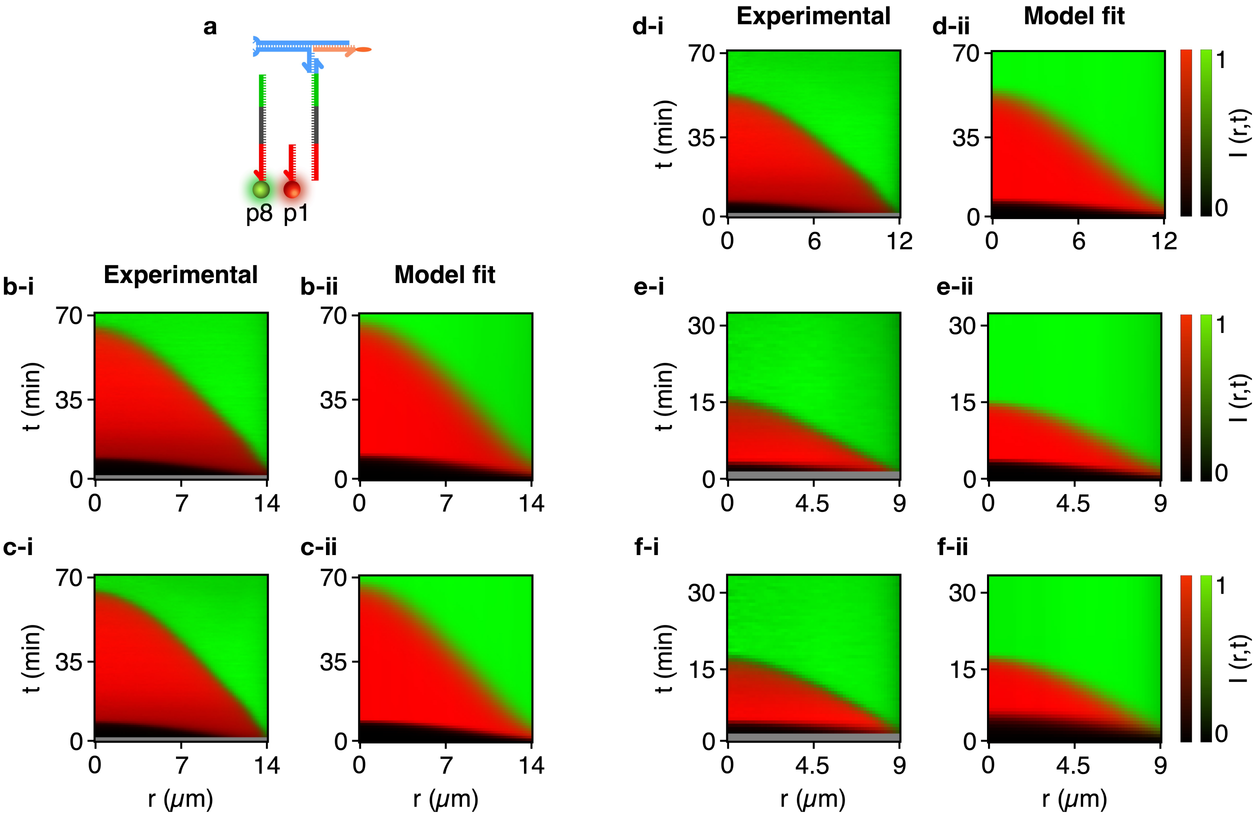

**Figure S5:** Experimental and modeled (fitted) results in 5 different DNA condensates of varying sizes showing the reaction diffusion behavior of two competing patterning strands, namely p1 and p8. **a**: Schematic depiction of the patterning and base strands. **b-f**: Experimental (**-i**) and modeled (**-ii**) color maps as used and discussed in Fig. 2. Gray bands in the experimental data mark the time gap between exposure of the condensates to patterning strands and the beginning of data acquisition (see Experimental Methods).

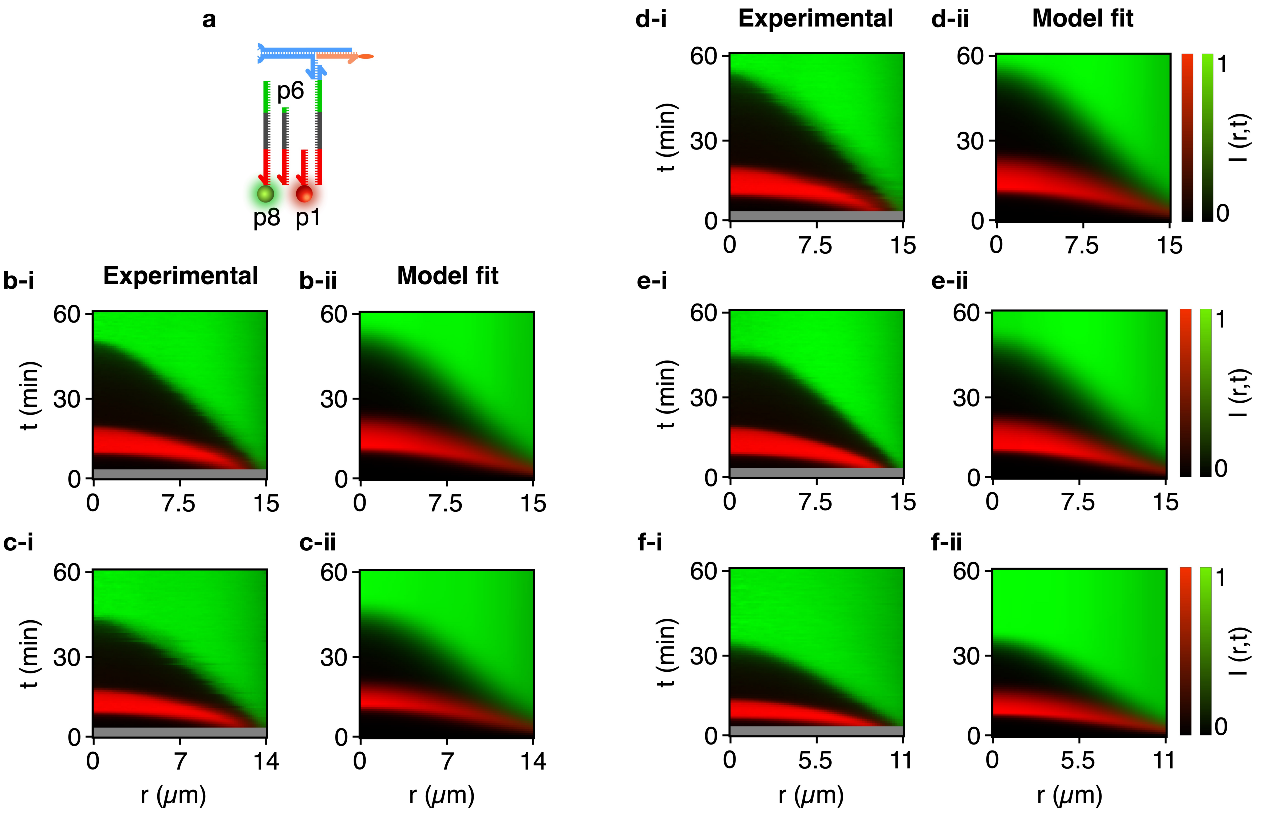

**Figure S6:** Experimental and modeled (fitted) results in 5 different DNA condensates of varying sizes showing the reaction diffusion behavior of three competing patterning strands, namely p1, p6 and p8. **a**: Schematic depiction of the patterning and base strands. **b-f**: Experimental (**-i**) and modeled (**-ii**) color maps as used and discussed in Fig. 2. Gray bands in the experimental data mark the time gap between exposure of the condensates to patterning strands and the beginning of data acquisition (see Experimental Methods).

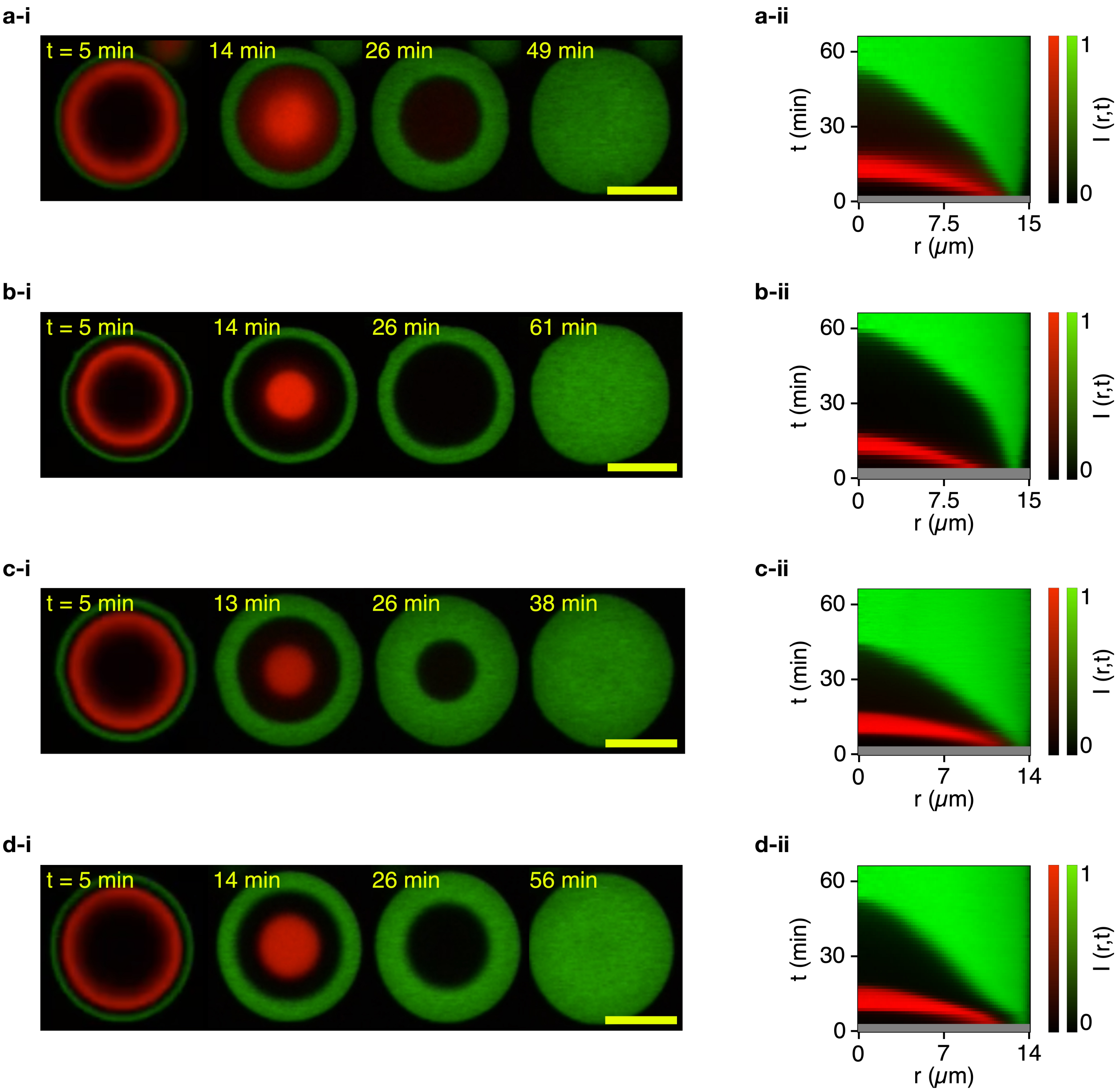

**Figure S7:** Control over the relative thickness of concentric domains can be achieved by tuning the length of patterning strands. Comparison of reaction diffusion behavior for different combinations of three patterning strands, in which the same longest and shortest strands are always used (p8 in green and p1 in red, respectively), while the intermediate length non-fluorescent strand is changed. **a**: p2, 20 nt. **b**: p4, 25 nt. **c**: p6, 30 nt. **d**: p7, 35 nt. The data is shown as confocal microscopy image series (**-i**) and micrographs (**-ii**). The scale bar corresponds to 15 $\mu$m

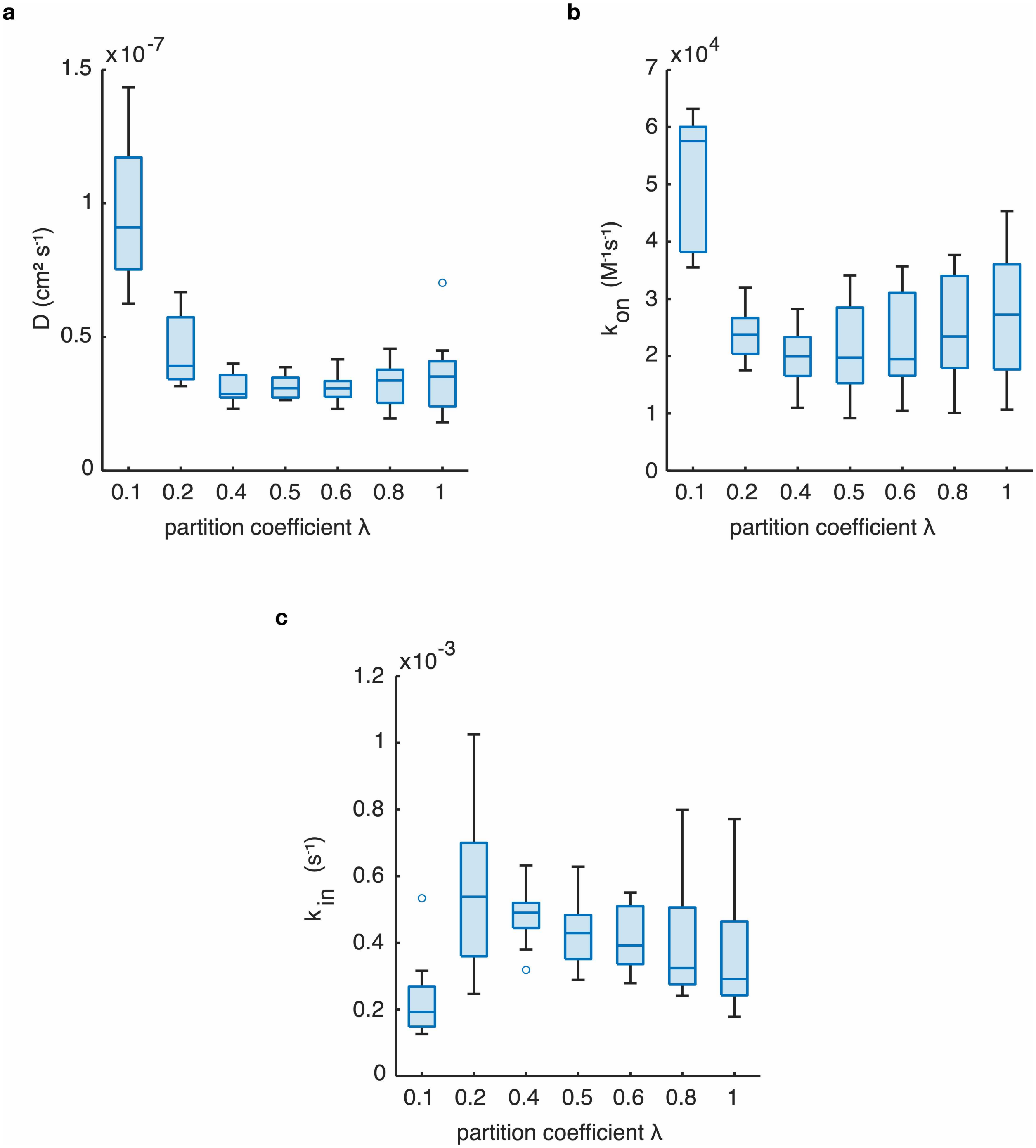

**Figure S8:** Effects of changing partition coefficient $\lambda$ (Eq. S7) on data fitting for the single patterning strand p8 (see Modeling Methods). **a**: Material exchange rate $k_{\mathrm{in}}$. **b**: Diffusion coefficient $D$. **c**: Binding rate constant $k_{\mathrm{on}}$. The partition coefficient is found to only affect the results when unrealistically small values are used, hence we set $\lambda = 1$ for our fitting.

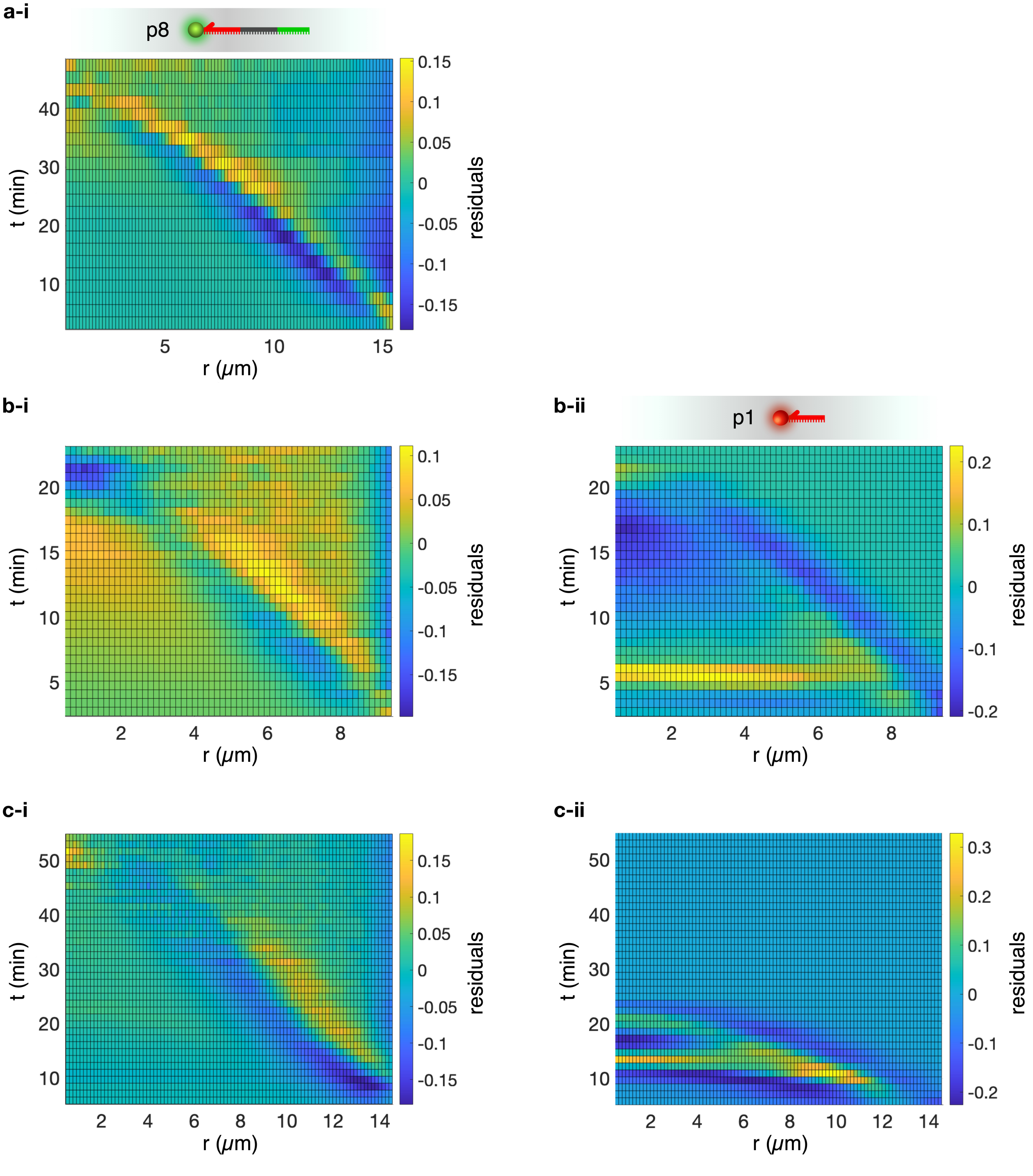

**Figure S9:** Fit residual maps showing $I_{\mathrm{exp}}(r,t) - I_{\mathrm{model}}(r,t)$, where $I_{\mathrm{exp}}(r,t)$ and $I_{\mathrm{model}}(r,t)$ are the experimental fluorescence intensity maps and the fitted outcome of the numerical model (see Experimental and Modeling Methods). Residual maps are shown for condensate-patterning experiments involving one, two and three patterning strands, as shown in Fig. 2**a**-**c**. Data are shown as a function of the distance from the center of the condensate, $r$, and the time elapsed from exposure of the condensates to the patterning strands, $t$. **a**: One green fluorescent (p8) patterning strand. **b**: Two patterning strands, green fluorescent (p8) and red fluorescent (p1). **c**: Three patterning strands, green fluorescent (p8), non-fluorescent (p6) and red fluorescent (p1). For panels **b** and **c**, residual plots are extracted for both fluorescent strands, p8 and p1, and shown in subplots **-i** and **-ii**, respectively. The emergence of patterns in the residual maps highlights small deviations between the experimental data and the fitting outcomes, expectedly localized around the propagation fronts.

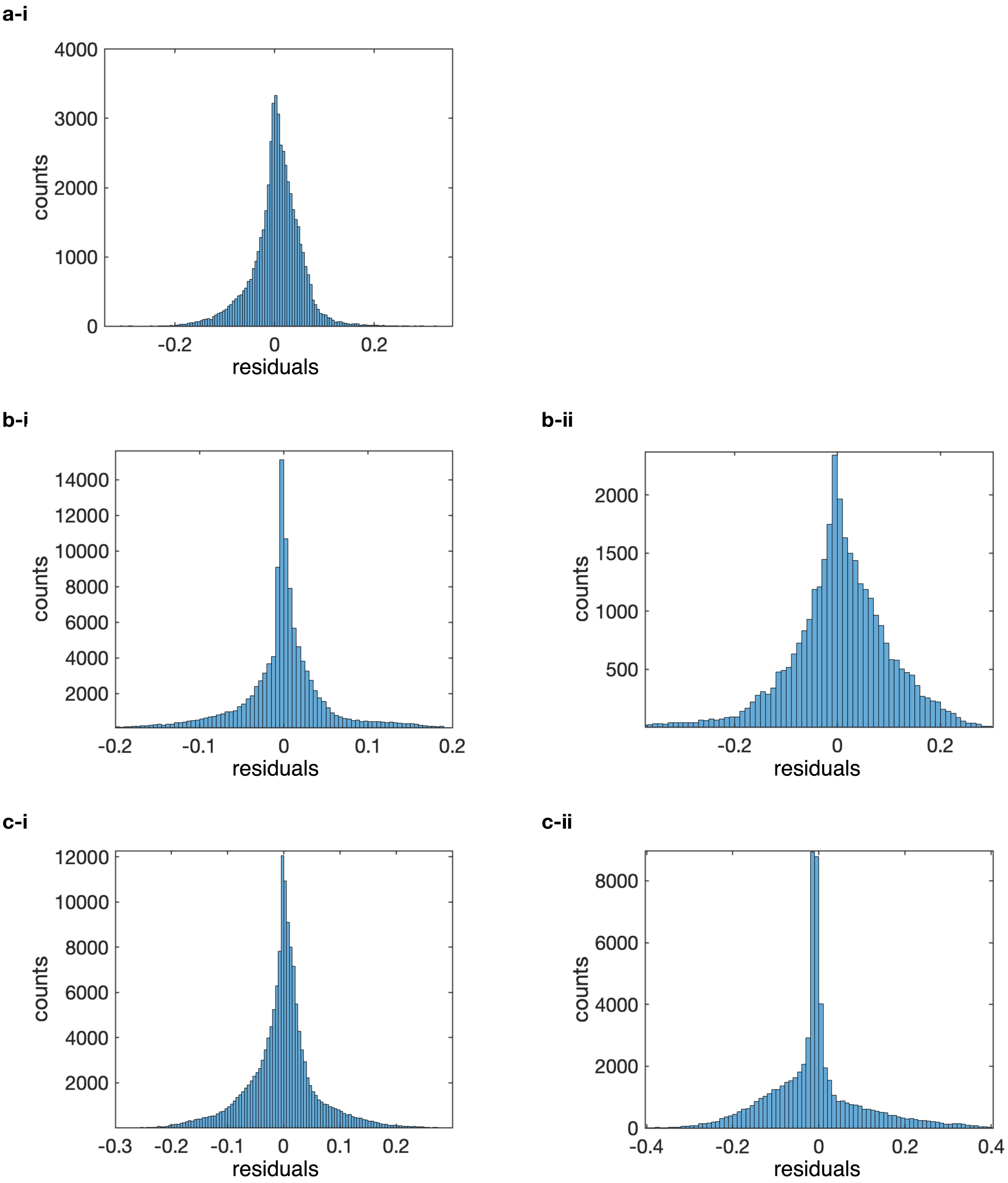

**Figure S10:** Histograms showing the distribution of the fitting residuals $I_{\mathrm{exp}}(r,t) - I_{\mathrm{model}}(r,t)$, where $I_{\mathrm{exp}}(r,t)$ and $I_{\mathrm{model}}(r,t)$ are the experimental fluorescence intensity maps and the fitted outcome of the numerical model (see Experimental and Modeling Methods). Data are shown for condensate-patterning experiments involving one, two and three patterning strands, as shown in Fig. 2**a-c**, collating residual values for all tested condensates for each condition. **a**: One green fluorescent (p8) patterning strand. **b**: Two patterning strands, green fluorescent (p8) and red fluorescent (p1). **c**: Three patterning strands, green fluorescent (p8), non-fluorescent (p6) and red fluorescent (p1). For panels **b** and **c**, residual plots are extracted for both fluorescent strands, p8 and p1, and shown in subplots **-i** and **-ii**, respectively.

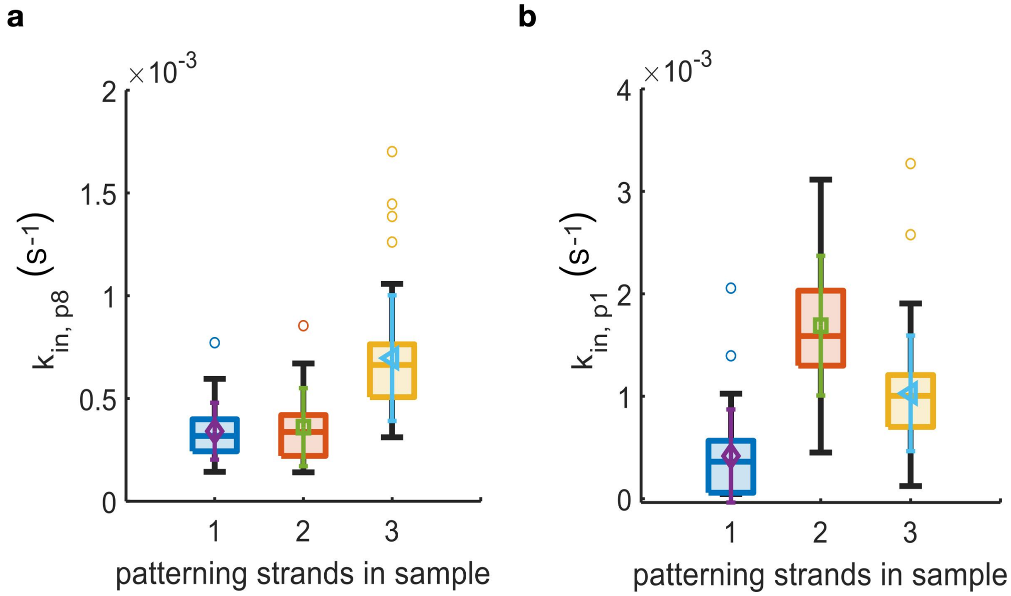

**Figure S11:** Rate of material exchange between condensate and bulk $k_{\mathrm{in}}$ (**a** and **b**, see Numerical Methods) as fitted for strands p8 (40 nt, **a**) and p1 (16 nt, **b**) in samples with a single patterning strand (p1 or p8; Fig. 2a and Figs S3-4; $N = 33$ condensates for p1 and $N = 29$ condensates for p8 ), two patterning strands (p1 and p8; Fig. 2b and Fig. S5; $N = 23$ condensates) and three patterning strands (p1, p6, and p8; Fig. 2c and Fig. S6; $N = 43$ condensates). The results are displayed as box plots with highlighted median, upper and lower quartiles (box), upper and lower 50th centiles (whiskers) and outliers excluded. Overlaid to the box plots are the mean (symbol) and standard deviation (same-color errorbar) of the distributions.

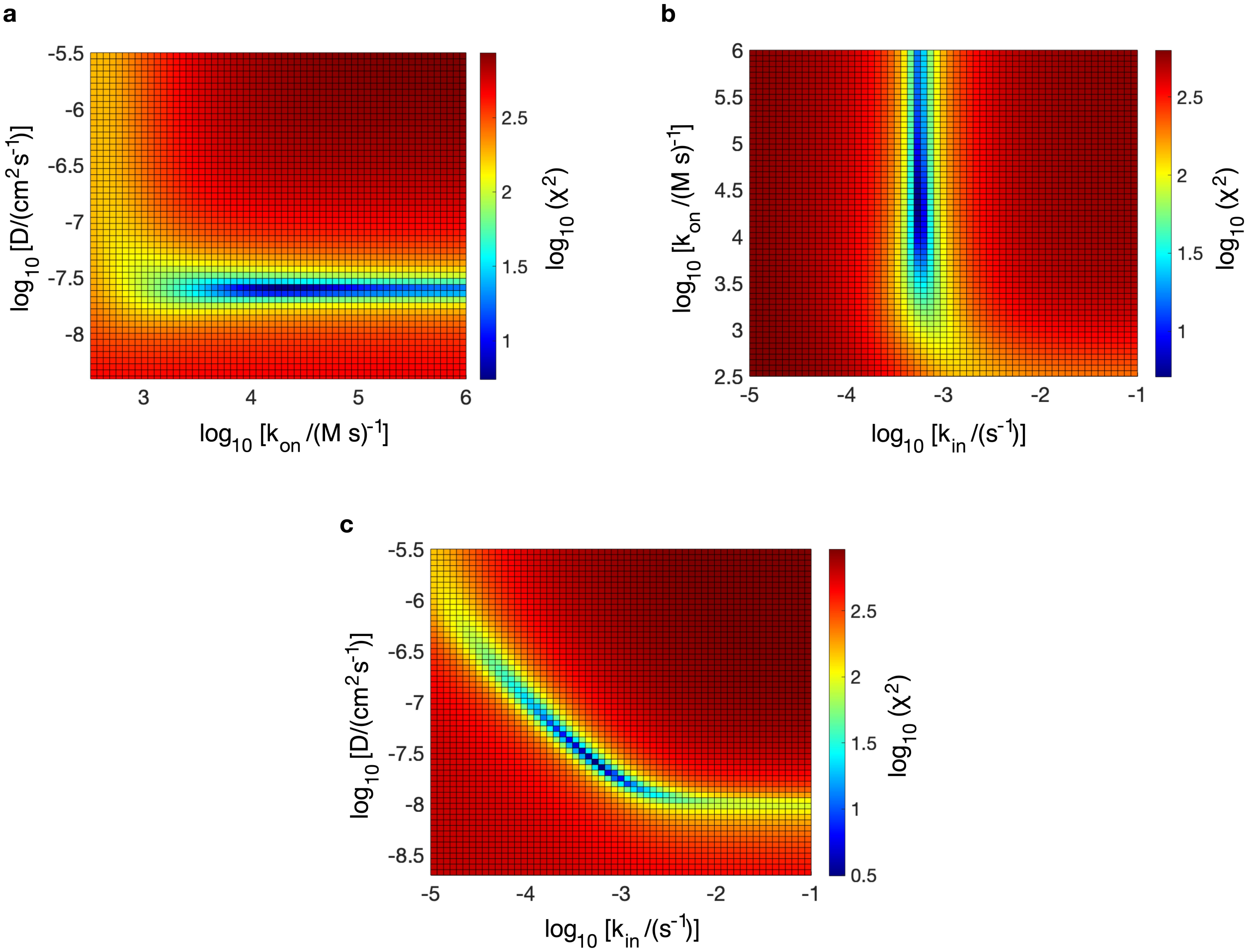

**Figure S12:** Heat maps of the sum of square residuals, $\chi^2$, shown as a function of pairs of fitting parameters appearing in the numerical reaction-diffusion model, namely the diffusion constant $D$, the binding rate $k_{\rm on}$ and the material exchange rate $k_{\rm in}$. $\chi^2$ is calculated as $\chi^2 = \sum_{r,t}([I_{\rm exp}(r,t) - I_{\rm model}(r,t)]^2)$, where $I_{\rm exp}(r,t)$ and $I_{\rm model}(r,t)$ are the experimental and modelled fluorescence intensity maps at (discretized) radial and time coordinates $r$ and $t$ (compare *e.g.* Fig. 2). **a:** $k_{\rm on}$ vs $D$. **b:** $k_{\rm in}$ vs $k_{\rm on}$. **c**: $k_{\rm in}$ vs $D$. Maps in **a** and **b** show that $k_{\rm on}$ is uncorrelated to both $D$ and $k_{\rm in}$. We note that $\chi^2$ increases slowly as $k_{\rm on}$ increases from its optimal value (minimum in $\chi^2$), indicating weak sensitivity of the model for $k_{\rm on} \gtrsim 10^5\,{\rm M}^{-1}\,{\rm s}^{-1}$. This feature is the result of the Gaussian blurring applied to the model output to emulate diffraction (see Modeling Methods), which has a similar effect on $I_{\rm model}(r,t)$ as decreasing $k_{\rm on}$. Panel **c** highlights a correlation between $D$ and $k_{\rm in}$ in the vicinity of the $\chi^2$ minimum. This is expected as the diffusion rate is dependent on $D$ and local molecular concentration (see Eq. S2), and the latter is affected by $k_{\rm in}$ through the boundary conditions (see Eq. S7). For all two-variable maps, the third parameter is kept constant at the value obtained from fitting the model to the experimental data. Data shown here correspond to a representative condensate patterned with a single p8 patterning strand (see Fig. 2**a**).

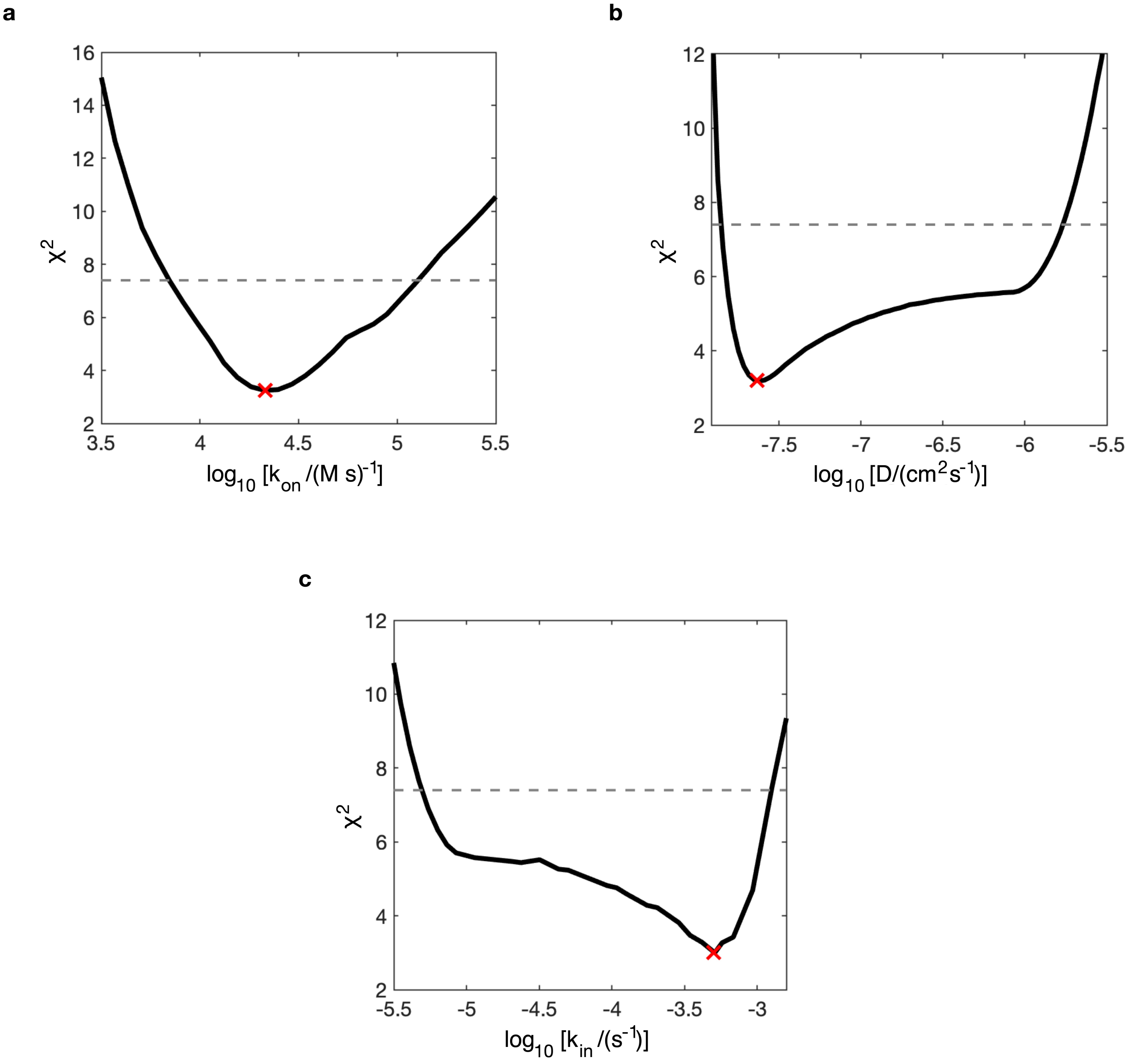

**Figure S13:** Profile likelihoods (black line) associated to the fit of an experimental fluorescence intensity map $I(r, t)$ to the output of our numerical reaction-diffusion model (see Experimental and Modeling Methods). Data are relative to a representative condensate patterned with a single p8 patterning strand (see Fig. 2**a**). Profiles are computed for each of the model parameters: $k_{\mathrm{on}}$ (**a**), $D$ (**b**) and $k_{\mathrm{in}}$ (**c**). To compute the likelihood curves, the parameter of interest (*e.g.* $k_{\mathrm{on}}$) is varied over the range of interest (2 to 2.5 orders of magnitude), and, for each selected value, the sum of square residuals $\chi^2$ is minimized against the other two parameters (see caption of Fig. S12 for definition of $\chi^2$). The red X shows optimal values of the relevant parameters as obtained by performing the fit, which, consistently, align to the minima of the likelihood curves. The dashed horizontal lines at $\chi^2 \simeq 7.4$ indicate the threshold for identifiability with 68.4% confidence, and are computed following references.[8–10] According to this threshold, all parameters are structurally identifiable with 68.4% confidence.[8–10] The intersection points between the dashed line and the solid line define the confidence intervals, which are $[6.982 \times 10^3,\ 1.275 \times 10^5]\,\mathrm{M^{-1}\,s^{-1}}$ for $k_{\mathrm{on}}$, $[1.404 \times 10^{-8},\ 1.705 \times 10^{-6}]\,\mathrm{cm^2\,s^{-1}}$ for $D$ and $[4.943 \times 10^{-6},\ 1.252 \times 10^{-3}]\,\mathrm{s^{-1}}$ for $k_{\mathrm{in}}$.

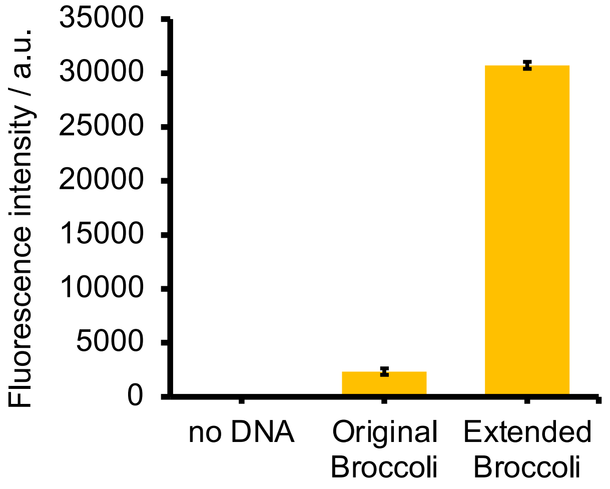

**Figure S14:** Increased fluorescence of an extended Broccoli RNA aptamer as determined with fluorimetry (see Experimental Methods). Error bars represent one standard deviation. See Table S1 for aptamer sequences.

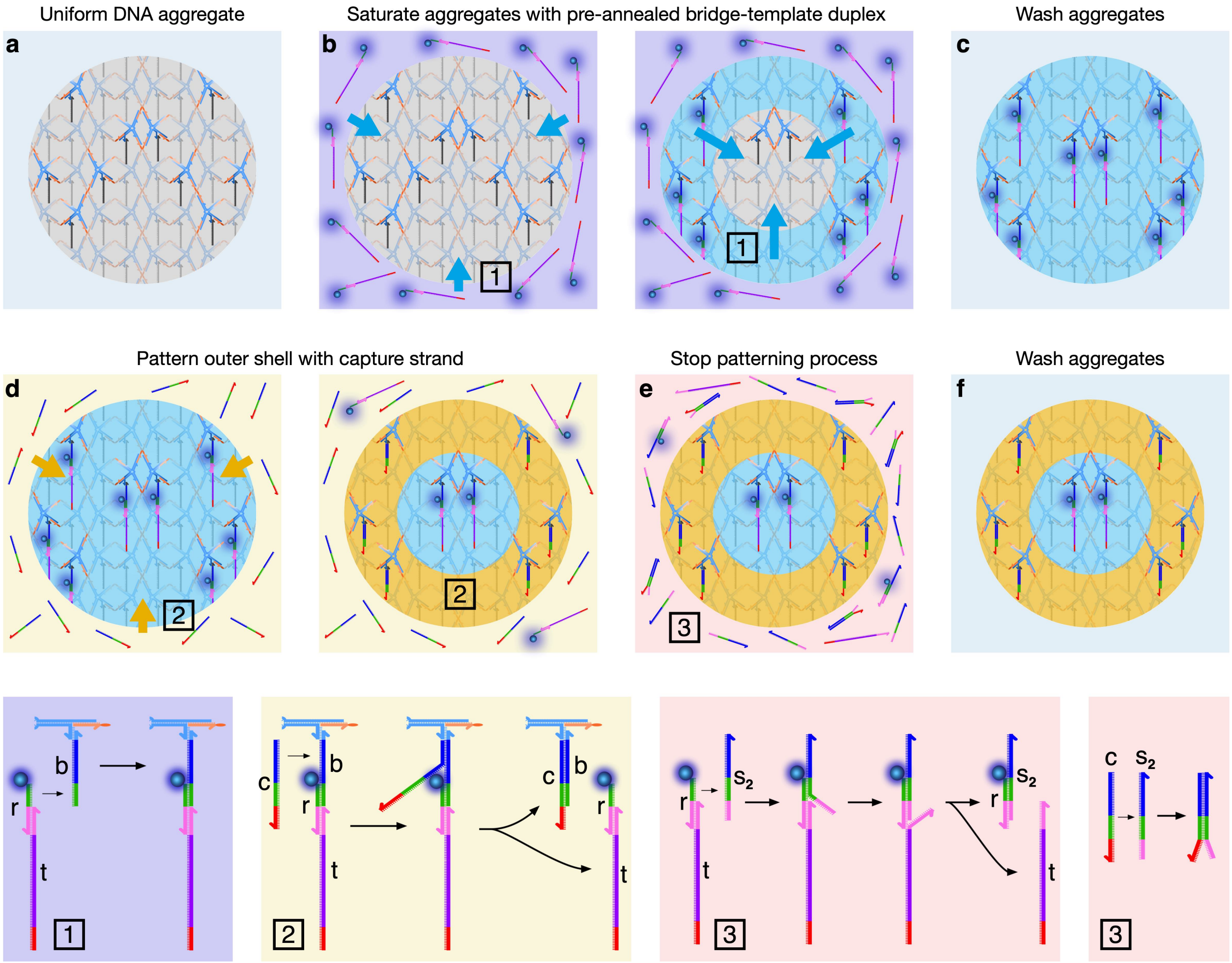

**Figure S15:** Nucleus-shell patterning process for RNA expression experiments. The aim is to connect two different complexes to the base strand (b), one in the core (nucleus) of a condensate and the other in the surrounding shell, thus creating two distinct environment with localized functionalities. In the nucleus, we aim to localize a *bridge* strand (r) and a *template* strand (t), which form a T7 promoter (pink) and a single stranded polymerase template. In the shell we localize a *capture* strand (c). Because the r-t duplex, which needs to be located in the nucleus, diffuses slower than c, a multi-step approach is required. **a:** The process starts with freshly extracted, washed condensates. **b:** An excess concentration of pre annealed r-t duplex is added to the sample and left to diffuse until all condensates are saturated. The detailed binding process is shown in highlight 1. **c:** Condensates are washed to remove excess r-t duplexes. **d:** Strand c is added to the aggregates and binds to strand b by displacing the r-t duplex. The duration of this process determines the width of the nucleus and shell. A longer wait leads to a thicker shell. The detailed displacement reaction is depicted in highlight 2. **e:** The patterning process of strand c is halted by adding the stop2 strand (s2) in excess. The stop2 strand sequesters all free c strands in solution and, in addition, it breaks apart any r-t construct in the sample that have been previously displaced by the c strands. The latter step is performed to ensure no RNA production from freely diffusing r-t constructs. Binding and displacing details are shown in highlight 3. Note how the s2-r construct does not feature a complete T7 promoter domain (pink), and thus is not able trigger transcription. **f:** As a final step, the condensates are further washed to remove any freely diffusing strands in the sample. All strand sequences are found in Table S1.

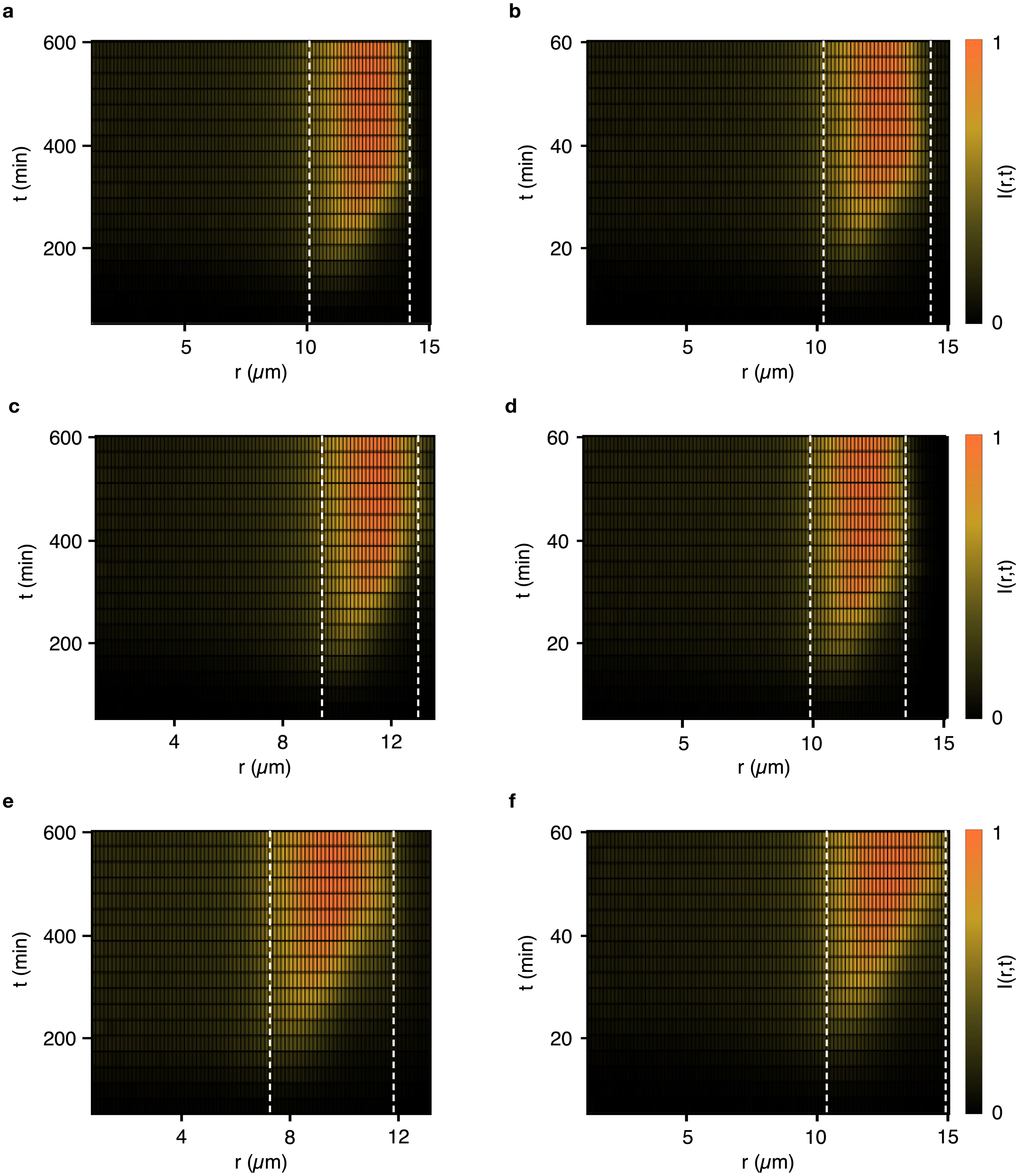

**Figure S16: a-f**: Time-evolution of the radial intensity profiles from the Broccoli aptamers produced in the nucleus of (various distinct) artificial cells, as they progressively accumulate in their shell, as discussed in Fig. 4. Note the intensity profile indicative of inside-out accumulation.

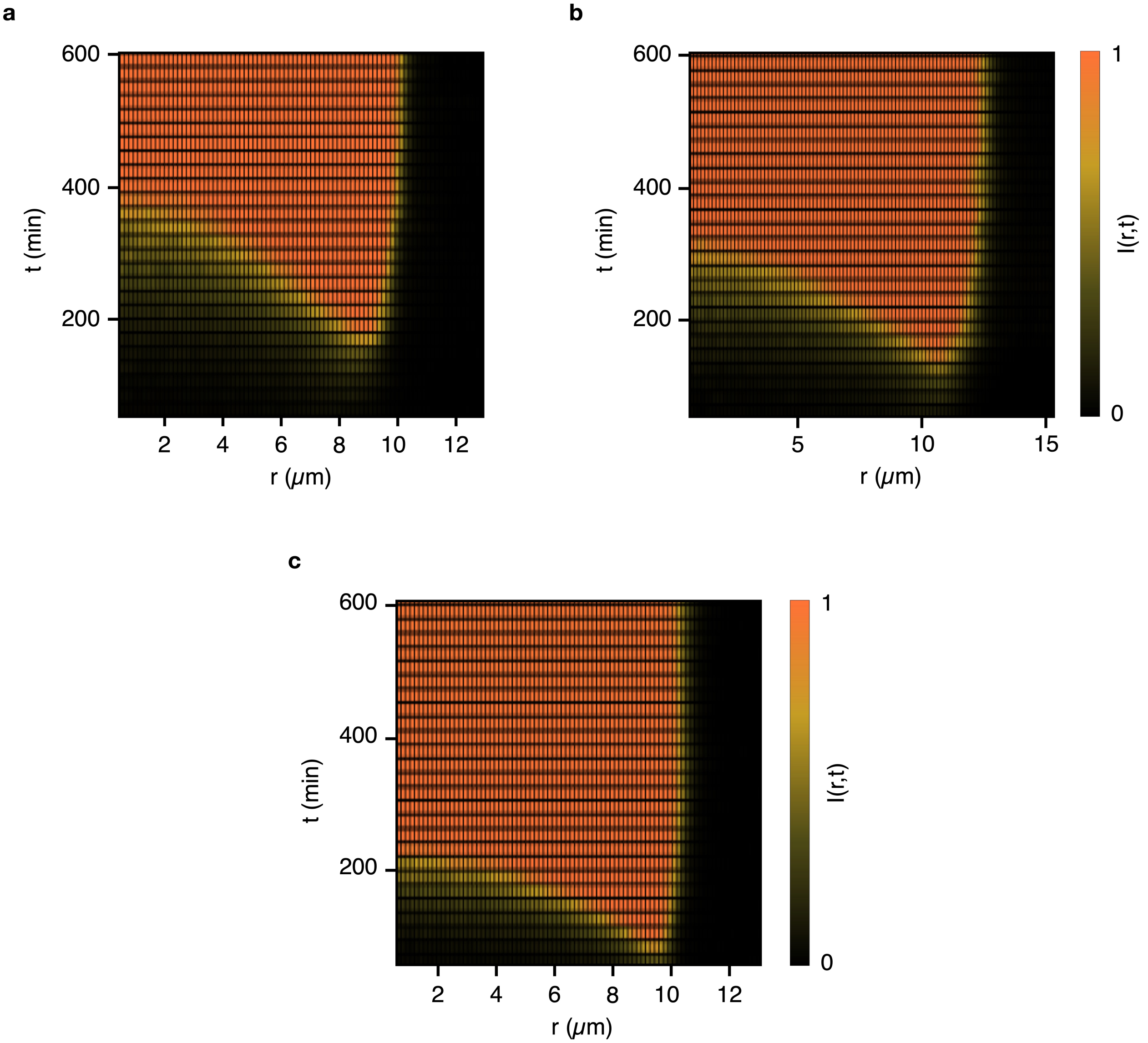

**Figure S17:** Broccoli aptamer synthesized in the bulk (outside the condensates) accumulates within the condensates from the outside-in. **a-c**: Color maps of time-evolution of the radial fluorescence intensity from Broccoli aptamer accumulating within condensates that feature capture strands but no template/bridge strands. Broccoli synthesis here occurs in the solution outside the condensates from freely diffusing bridge-template duplexes. Note the clear difference in the patterns compared with the artificial cells in Fig. 4 and Fig. S16, where the aptamer is synthesized within the nucleus and accumulates within the shell from the inside-out. The concentration of freely diffusing bridge-template duplexes for the different samples are 0.15 µM in **a**, 0.45 µM in **b** and 0.9 µM in **c**.

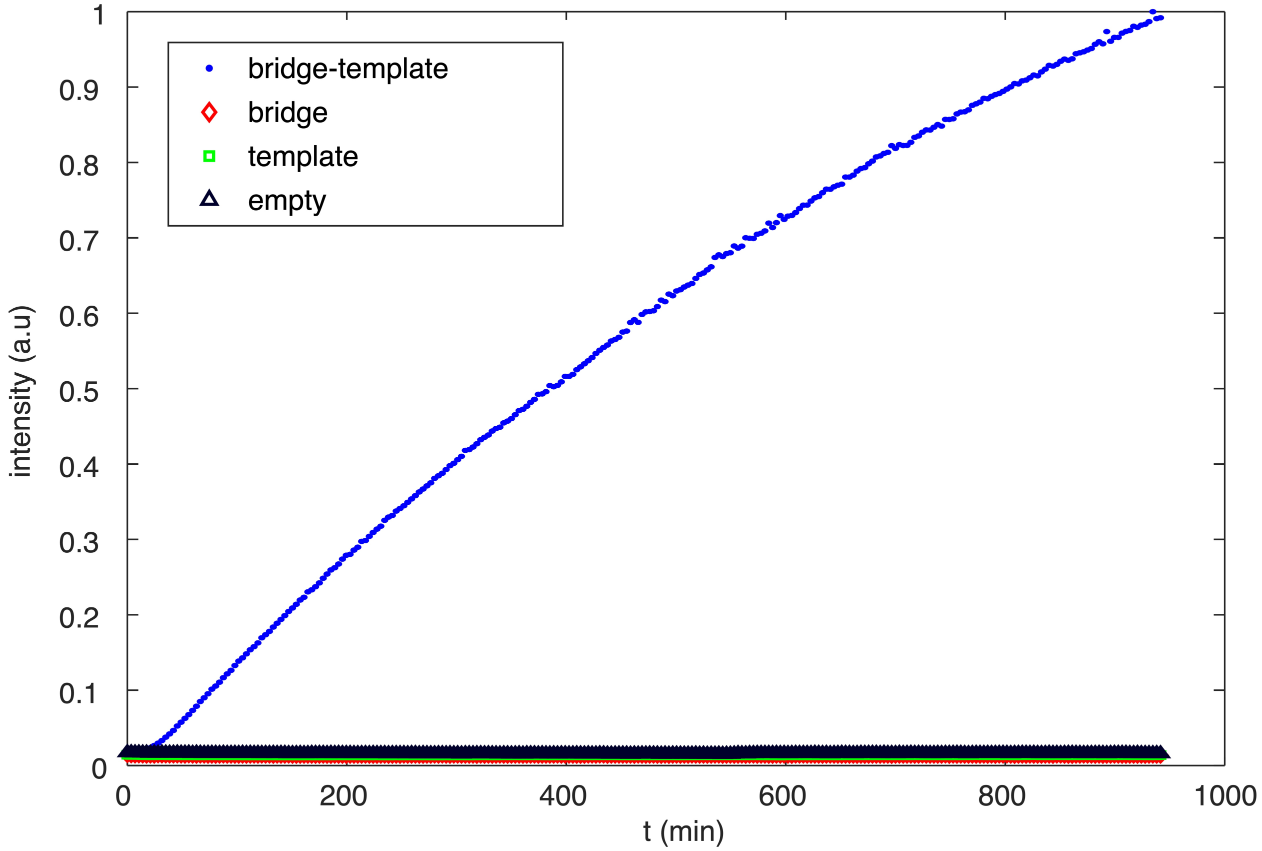

**Figure S18:** Bulk fluorimetry control experiment demonstrating that the fluorescent Broccoli aptamer is only synthesized in the presence of both the template strands and the bridge strand that hybridizing with the template strand forms the T7 promoter region. Here all components are freely diffusing rather than bound within condensates. See Table S1 for strand sequences.

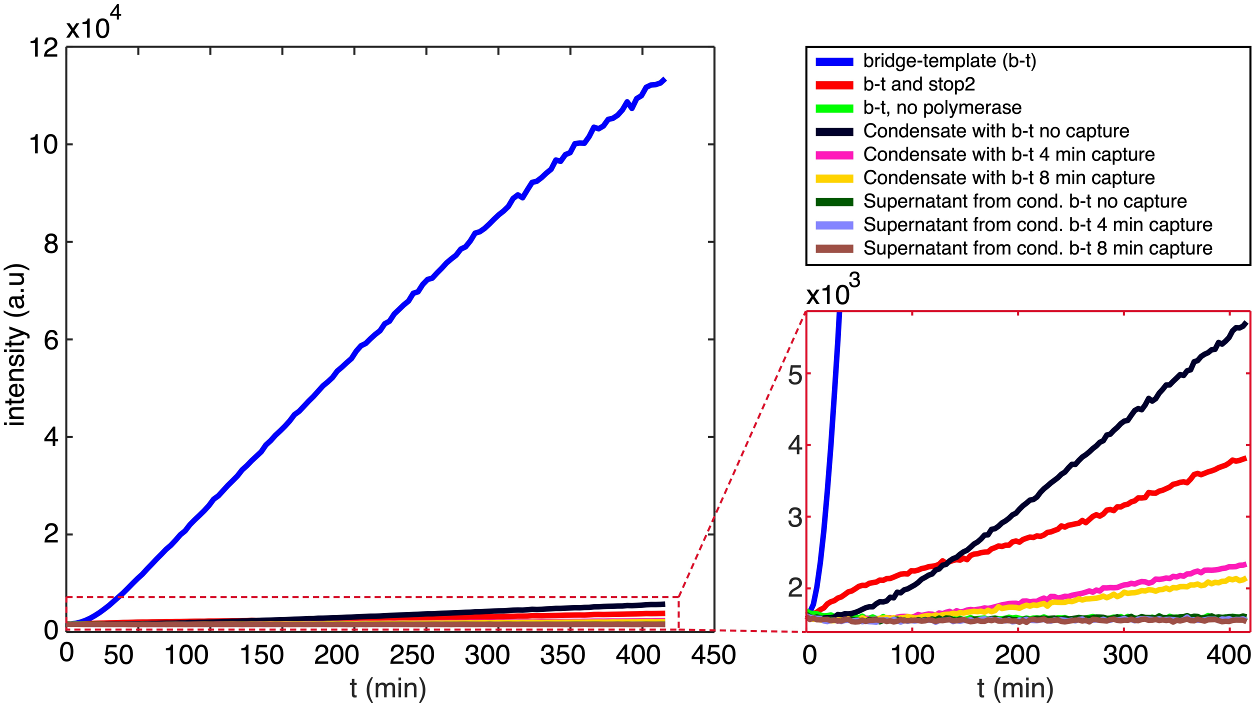

**Figure S19:** Bulk fluorimetry control experiment monitoring Broccoli synthesis in various conditions, including freely diffusing bridge-template (b-t) constructs (with and without the addition of the stop2 strand), condensates (ACs) hosting b-t constructs, and supernatants from the condensate washing steps (see Experimental Methods). The fluorescent signal from freely-diffusing b-t constructs (at concentration of 0.5 µM) is strongly suppressed when the stop2 strand is added (at concentration of 2 µM), confirming its ability to inhibit broccoli synthesis by disrupting the promoter region in the b-t construct through toehold-mediated strand displacement. Curves relative to condensates are substantially dimmer compared to data collected in bulk, likely as a result of the overall smaller amount of b-t construct present and/or reduced activity of the T7 polymerase within the condensates. Note that condensates patterned with the capture constructs for increasing amounts of time produce weaker signals, as expected given that the size of the Broccoli-producing nucleus progressively shrinks (see Experimental Methods). No signal above background (b-t, no polymerase) is recorded in the supernatants collected at the final washing step in the artificial cell preparation protocol (see Experimental Methods), confirming that Broccoli synthesis demonstrated in Figs. 4 and S16 occurs uniquely within the nucleus of the artificial cells. See Table S1 for strand sequences.

### Supplementary tables

**Table S1: Oligonucleotide sequences.** All strands and sequences used to make condensates, reaction diffusion patterning, and polymerase experiments in 5' to 3' direction.

| Strand names | Sequences (5' → 3') |
|---|---|
| core1 | CGACGCCGTGACGCCGTGGCCTGTGATTGAGGCGCTGCGTCGTCCACCGTGT-GAAACTTGTCCGTTCTAAATC |
| core2 | CGACGCCGTGACGCGTTTCACACGGTGGACGACGCTCGGACTAGAACTGTCTCGAAC |
| core3 | CGACGCCGTGACGCGTTCGAGACAGTTCTAGTCCGTCGCGAATACGCCGTGCCGTGC |
| core4 | CGACGCCGTGACGCGCACGGCACGGCGTATTCGCGTGCGCCTCAATCACAGGCCACG |
| cholesterol | GCGTCACGGCGTCGAA TEG Cholesterol |
| base (b) | GGTGAGGTGAGTGGAGTGGAGGTGTAGGAGTGAGGGTAAGGATTTAGAACGGAC |
| p1 (16nt) | CCACTCACCTCACCTT Alexa 594 |
| p2 (20nt) | TCCACTCCACTCACCTCACC |
| p3 (22nt) | CCTCCACTCCACTCACCTCACC |
| p4 (25nt) | ACACCTCCACTCCACTCACCTCACC |
| p5 (28nt) | CCTACACCTCCACTCCACTCACCTCACC Alexa 488 |
| p6 (30nt) | CTCCTACACCTCCACTCCACTCACCTCACC |
| p7 (35nt) | CCTCACTCCTACACCTCCACTCCACTCACCTCACC |
| p8 (40nt) | CTTACCCTCACTCCTACACCTCCACTCCACTCACCTCACC Alexa 488 |
| stop (s) | GGTGAGGTGAGTGGAGTGGAGGTGTAGGAGTGAGGGTAAG |
| bridge (r) | Alexa 647N CCACTCACCTCACCTAATACGACTCACTATA |
| template (t) | GACCTAGACCGAACGGGTCTAGGAGCCCACACTCTACTCGACAGATACGAATATCTGGAC-CCGACCGTCTCCTAGACCCTATAGTGAGTCGTATTA |
| Original Broccoli | AUUAUGCUGAGUGAUAUCCCAGCUCUGCCAGCCCAGGUCUAUAAGCAUAGACAGCUCAU-CUCACACCCGAG |
| Extended Broccoli | GGGUCUAGGAGACGGUCGGGUCCAGAUAUUCGUAUCUGUCGAGUAGAGUGUGGGCUCCUA-GACCCGUUCGGUCUAGGUC |
| capture (c) | CTTACCCTCACTCCTACACCTCCACTCCACTCACCTCACCGACCTAGACCGAAC |
| stop2 | TGAGTCGTATTAGGTGAGGTGAGTGGAGTGGAGGTGTAGGAGTGAGGGTAAG |

#### Supplementary videos key

**Video S1**: 1 patterning strand (p8) – large view

**Video S2**: 1 patterning strand (p8) – zoomed-in view

**Video S3**: 2 patterning strands (p1 and p8) – large view

**Video S4**: 2 patterning strands (p1 and p8) – zoomed-in view

**Video S5**: 3 patterning strands (p1, p6 and p8) – large view

**Video S6**: 3 patterning strands (p1, p6 and p8) – zoomed-in view

**Video S7**: 5 patterning strands (p1, p3, p5, p7 and p8) – large view

**Video S8**: 5 patterning strands (p1, p3, p5, p7 and p8) – zoomed-in view

**Video S9**: 3 patterning strands (p1, p6 and p8) and stop strand – large view

**Video S10**: 3 patterning strands (p1, p6 and p8) and stop strand – zoomed-in view

**Video S11:** 1 patterning strand (p1) – large view

**Video S12**: 1 patterning strand (p1) – zoomed-in view

**Video S13**: Broccoli RNA production and storage in artificial cells – example 1

**Video S14**: Broccoli RNA production and storage in artificial cells – example 2

**Video S15**: Broccoli RNA production and storage in artificial cells – zoomed-in view (fluorescence)

**Video S16**: Broccoli RNA production and storage in artificial cells – zoomed-in view (brightfield + light fluorescence)
